# Fibre-specific laterality of white matter in left and right language dominant people

**DOI:** 10.1101/2020.08.04.236000

**Authors:** Helena Verhelst, Thijs Dhollander, Robin Gerrits, Guy Vingerhoets

## Abstract

Despite the typical symmetrical appearance of the human brain, several functional and structural asymmetries have been reported. Language is the most commonly described lateralised cognitive function, relying relatively more on the left hemisphere in over 90% of the population. This is in line with white matter studies which have revealed leftwards lateralisation of the arcuate fasciculus, a white matter tract that connects important language-related regions. Most research to date examining the structure-function relationship of language lateralisation only included people showing a left language hemisphere dominance. As such, the reported correlations do not allow for inferences of relationships between the directions of functional and structural lateralisation of language.

In this work, we applied a state-of-the-art “fixel-based” analysis approach, which allows to statistically analyse white matter micro- and macrostructure on a fibre-specific level. To study lateralisation using this framework, we defined a bespoke fibre-specific laterality index which allowed us to examine whole-brain white matter asymmetries in samples of participants with left and right language dominance (LLD and RLD respectively). Both groups showed similar extensive and intricate patterns of significant white matter lateralisation. Few group differences were found between both groups, with a similar leftwards lateralisation of the arcuate fasciculus, regardless of functional language lateralisation of the participants. A significant group difference of lateralisation was detected in the forceps minor, with a leftwards lateralisation in LLD and rightwards lateralisation for the RLD group.

In conclusion, we showed that fixel-based analysis of fibre-specific lateralisation indices is an effective approach to study white matter asymmetries. Our results suggest that the lateralisation of language functioning and the arcuate fasciculus are driven by independent biases. The exact relationship between forceps minor asymmetry and language dominance could be an interesting subject of future studies.

## Introduction

Although the human brain may appear to be a symmetrical organ at first sight, ample studies have revealed specific differences between the left and right hemispheres, both functionally and structurally (Güntürkün & Ocklenburg, 2017). Regarding functional lateralisation, there is broad consensus on which functions typically lateralise to which hemisphere (Vingerhoets, 2019). For example, one of the best documented and robust population-based biases relates to language function, which is predominantly driven by the left hemisphere in well over 90% of the population (Mazoyer et al., 2014; Van der Haegen et al., 2013; Westerhausen et al., 2014). Regarding structural asymmetry, however, the literature is more ambiguous. The most studied white matter tract with respect to lateralisation is the arcuate fasciculus, a perisylvian pathway connecting important language related regions, including Broca’s and Wernicke’s areas (Catani et al., 2005). Several studies have shown a leftwards lateralisation of this white matter bundle, on both the macro- and microscopic scale (Bain et al., 2019; Catani et al., 2007; Slater et al., 2019). The lateralisation patterns for other tracts are less examined and reported results are sometimes inconsistent. The uncinate, for example, has been found to be rightwards lateralised by some studies (Highley et al., 2002; Park et al., 2004), while others report a leftwards lateralisation (Ocklenburg et al., 2013; Slater et al., 2019) or an absence of structural asymmetry (Mannaert et al., 2019; Thiebaut de Schotten, ffytche, et al., 2011).

White matter structure likely plays a role in the development of functional lateralisation as it links different grey matter regions responsible for cognitive functioning (Ocklenburg et al., 2016). Several studies have reported on the relationship between the degree of functional and structural lateralisation of cognitive functioning. Thiebaut de Schotten and colleagues, for example, found a correlation between diffusion MRI-derived metrics of lateralisation of the superior longitudinal fasciculus and visuospatial functioning (Thiebaut de Schotten, Dell’Acqua, et al., 2011). A correlation between brain activation during language-related tasks and structural asymmetries in white matter has been reported in multiple studies; in children, as well as healthy adults and clinical groups (for a review see Ocklenburg et al. (2016)). However, most of these studies were performed in right-handers, who only rarely demonstrate rightward language laterality. As such, the reported correlations do not allow to conclude that the direction of functional lateralisation is related to the direction of structural lateralisation of language. Only a single study, by Vernooij et al. (2007), examined white matter asymmetries in a sample which included five left-handed participants with a right hemispheric language dominance. Regardless of the direction of functional language lateralisation, a leftwards asymmetry in relative streamline count (as derived from fibre tractography) of the arcuate fasciculus was reported. The results of Vernooij and colleagues suggest that structural asymmetry of the arcuate does not necessarily reflect functional hemispheric language lateralisation as previously hypothesized in the other studies. However, it has since been established that streamline counts from fibre tractography do not represent an appropriate metric to quantify white matter connectivity (Jones et al., 2013).

Most work on structural lateralisation to date has relied on diffusion tensor imaging (DTI) derived metrics, like fractional anisotropy (FA) or mean diffusivity (MD). It has been shown that DTI-based metrics are not capable of capturing multiple fibre orientations within a voxel—often referred to as “crossing fibres”—which appear to be very prevalent across the white matter (Jeurissen et al., 2013). Additionally, while DTI metrics have proven to be sensitive to certain white matter changes, they are nonspecific to changes in properties of individual fibre populations, often leading to non-intuitive effects and misleading interpretation in the presence of crossing fibres (Mito et al., 2018). To overcome these limitations, the fixel-based analysis (FBA) framework was recently proposed. FBA enables the quantification of micro- and macrostructural properties that are associated with specific ***fi***bre populations within a vo***xel*** (named “fixels”) (Raffelt et al., 2017). FBA studies typically achieve this using three metrics: fibre density (FD), fibre-bundle cross-section (FC) and fibre density and cross-section (FDC). FD is proportional to the total intra-axonal volume of axons along the orientation of a fibre population in each voxel (i.e. a microstructural measure, related to how densely axons are packed), whereas FC is proportional to the cross-sectional area of the white matter bundle in the region of each fixel (i.e. a macrostructural measure). FDC, defined simply as the product of FD and FC (i.e. the local density times the overall size), is thus sensitised to overall changes to the presence of axonal matter for individual fibre populations in each voxel. Even though FBA is a relatively recently developed method, it has proven to be a favourable technique to examine white matter properties in several studies. It has already been used in a range of different clinical groups (including Parkinson’s disease (Rau et al., 2019), traumatic brain injury (Verhelst et al., 2019), stroke (Egorova et al., 2020), multiple sclerosis (Gajamange et al., 2018), depression (Lyon et al., 2019), autism spectrum disorder (Dimond et al., 2019), and Alzheimer’s disease (Mito et al., 2018)) and in healthy cohorts (Choy et al., 2020; Dimond et al., 2020; Genc et al., 2018).

In the current study we aimed to examine structural white matter asymmetries in an unprecedented sample of 23 left-handers with right language dominance (RLD), complemented with a group of 38 individuals with left language dominance (LLD), to extend previous research on the structure-function relationship of language. We applied a state-of-the-art FBA approach and defined a bespoke fixel-wise asymmetry measure or laterality index (LI) designed for the FD, FC and FDC metrics. This enabled us to quantify and detect significant asymmetries in white matter micro- and macrostructure across all white matter for participants with left and right language dominance, as well as to quantify and detect significant relative differences of lateralisation between these groups.

## Methods

### Participants

As people with a right (hemisphere) language dominance (RLD) are uncommon, a specific recruitment strategy was pursued. Individuals with suspected RLD (based on a behavioural visual half field test) were invited to participate in the study. With the aim of increasing the likelihood of identifying people with RLD, we limited the inclusion for our study to left-handers only as it has been shown that RLD is more likely to occur in left-handers (15% to 34% of left-handers compared to only 4% to 14% of right-handers (Carey & Johnstone, 2014; Knecht et al., 2000)). Details of the recruitment procedure and the classification of the participants as RLD or left language dominance (LLD) based on an fMRI word generation task are documented elsewhere (Gerrits et al., 2020). The sample consisted of 24 participants with RLD and 39 with LLD. One LLD and one RLD participant were excluded, respectively due to a callosal abnormality visible on the T1w and diffusion MRI (dMRI) scans and imaging artefacts. This resulted in a final sample of 23 RLD and 38 LLD participants.

The LLD and RLD groups did not differ significantly in terms of age (LLD: 20 ± 3.98 years, RLD: 20 ± 3.50 years, U = 463.50, p = 0.69), sex distribution (LLD: 89% female, RLD: 87% female, χ^2^(1) = 0.09, p = 0.77), or years of formal education (LLD: 14 ± 1.66 years, RLD: 14 ± 1.56 years, U = 446.50, p = 0.89). All participants had normal or corrected-to-normal vision and reported no history of brain injury or developmental disorders. The study was approved by the medical ethical committee of Ghent University Hospital. Written informed consent was obtained from each participant.

### MRI data acquisition

MRI data were acquired using a 3 T Siemens PRISMA scanner with a 64-channel head coil. Diffusion MRI (dMRI) data was acquired for 6, 23, 42 and 70 non-collinear gradient directions respectively using b-values of 0, 700, 1200 and 2800 s/mm^2^. Additionally, an extra pair of b = 0 images was acquired using opposite phase encodings, to allow for correction of susceptibility induced EPI distortions. Other acquisition parameters are as follows: 2 mm isotropic voxel size, 75 slices, FoV = 240 mm, TR/TE = 3300 ms/76 ms.

### Fixel-based analyses of laterality

#### Fixel-based analysis processing pipeline

A fixel-based analysis (FBA) approach was used to enable fixel-wise quantification and statistical testing of fibre-specific laterality of white matter. FBA was implemented using MRtrix3 (Tournier et al., 2019) in accordance with the recommended procedures (Raffelt et al., 2017) and adapted where necessary to permit the calculation of bespoke laterality indices (LIs) for the traditional FBA metrics.

Preprocessing of the dMRI data included denoising (Veraart et al., 2016), removal of Gibbs ringing artefacts (Kellner et al., 2016), and motion, eddy-current, and EPI distortion corrections (Andersson et al., 2003, 2016; Andersson & Sotiropoulos, 2016). After upsampling the preprocessed data to a 1.5 mm isotropic voxel size, the white matter fibre orientation distributions (FODs) were estimated by a 3-tissue constrained spherical deconvolution approach (Jeurissen et al., 2014) using the group averaged tissue response functions for white matter, grey matter and cerebrospinal fluid (Dhollander et al., 2016, 2019). Multi-tissue informed log-domain intensity normalisation was performed to correct for bias fields and global intensity differences between participants.

To determine the homologous fixel pairs (i.e. corresponding left and right hemisphere fixels), additional “mirrored” FOD images were created for all participants by flipping the image data along the left-right axis (including similarly flipping the FOD in each voxel). Next, a study-specific unbiased and *symmetric* FOD template was generated based on a subset of the participants, using a similar approach to previous studies (Arun et al., 2019; Raffelt et al., 2017). To this end, 30 individuals (15 RLD and 15 LLD) were randomly selected. The symmetric template was then constructed by registering the 60 FOD images (both 30 “original” and 30 flipped) using an iterative nonlinear registration and averaging approach, which includes FOD reorientation (Raffelt et al., 2011). By registering both the original and flipped FOD images of each participant to the symmetric group template, FODs in the right hemisphere of the original FOD image were mapped to the corresponding FODs in the left hemisphere of the flipped FOD image (and vice versa). Next, a fixel analysis mask was defined by segmenting the individual fixels from the symmetric population FOD template subject to an FOD amplitude threshold of 0.9 (Raffelt et al., 2017). Similarly, fixels were also segmented from the individual subject FOD images (both “original” and flipped) in template space. Finally, correspondence was established between each fixel in the fixel analysis mask and a matching fixel in each individual subject FOD image (“original” and flipped) in template space. The resulting corresponding fixels of an individual subject are subsequently considered homologous fixel pairs. This then allows to compute laterality indices using metrics of each fixel pair, as well as to compare such laterality indices across different participants, whose homologous fixel pairs are similarly mapped to the fixel analysis mask.

The three traditional fixel-wise FBA metrics were then computed for all participants. The apparent fibre density (FD) was computed as the integral of the FOD lobe corresponding to each fixel. The fibre-bundle cross-section (FC) metric was computed based on the warps that were generated during registration to the symmetric FOD template, as the Jacobian determinant in the plane perpendicular to the fixel orientation (Raffelt et al., 2017). Finally, the combined measure of fibre density and cross-section (FDC) was calculated as the product of FD and FC.

#### Fixel-wise laterality index and fixel analysis mask

For each homologous fixel pair (i.e. right and corresponding left hemisphere fixel), we defined a bespoke laterality index (LI) for each FBA metric, as the (natural) log-ratio of the right and left hemisphere metric values. For example, the fixel-wise LI(FD) is defined as:

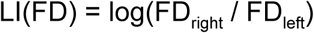

Both LI(FC) and LI(FDC) are defined analogously. An LI value of 0 indicates absence of lateralisation, whereas a positive LI value quantifies the degree of rightwards lateralisation and a negative LI value the degree of leftwards lateralisation. The log-ratio is the only measure of *relative* difference featuring the combination of three favourable properties: it is normed, symmetric and additive (Tornqvist et al., 1985). This differs from the somewhat “traditional” index of relative difference often used in lateralisation research (e.g. for functional MRI data), which is computed as the difference divided by the sum, sometimes multiplied by a constant (Seghier, 2008). The latter is, however, bounded and therefore lacks the additivity property. In Appendix A, we provide a brief comparison of both metrics for intuition as to how they relate to each other.

In practice, we computed the LI value from the corresponding fixel-wise values of the “original” and flipped subject images, e.g., log(FD_original_ / FD_flipped_). Hence, in the right hemisphere of the fixel analysis mask, this evaluates to the LI value as defined above. The left hemisphere of the fixel analysis mask therefore serves no additional purpose to our analysis (it represents the negative LI value, due to symmetry of both template and LI). Finally, a practical challenge introduced by the FBA framework lies in the fact that correspondence between some fixels of the analysis mask and those of individual participants sometimes cannot be established. While this poses no problems for the FC metric (which can *always* be derived from the subject-to-template warp combined with the fixels in the analysis mask), the FD value in this case is typically set to zero (Raffelt et al., 2017). This might introduce biases in FBA in general, but poses a unique challenge for the log-ratio in our definition of LI, which then cannot be evaluated. In this work, we mitigate this problem by setting the FD to a non-zero low value of 0.01 instead (this equals 1% of one calibrated unit of FD). Additionally, we further restrict the fixel analysis mask to only contain fixels for which correspondence with at least 90% of all participant hemisphere data could be established. Part of the final fixel analysis mask is shown in Figure 1.

**Figure 1:**
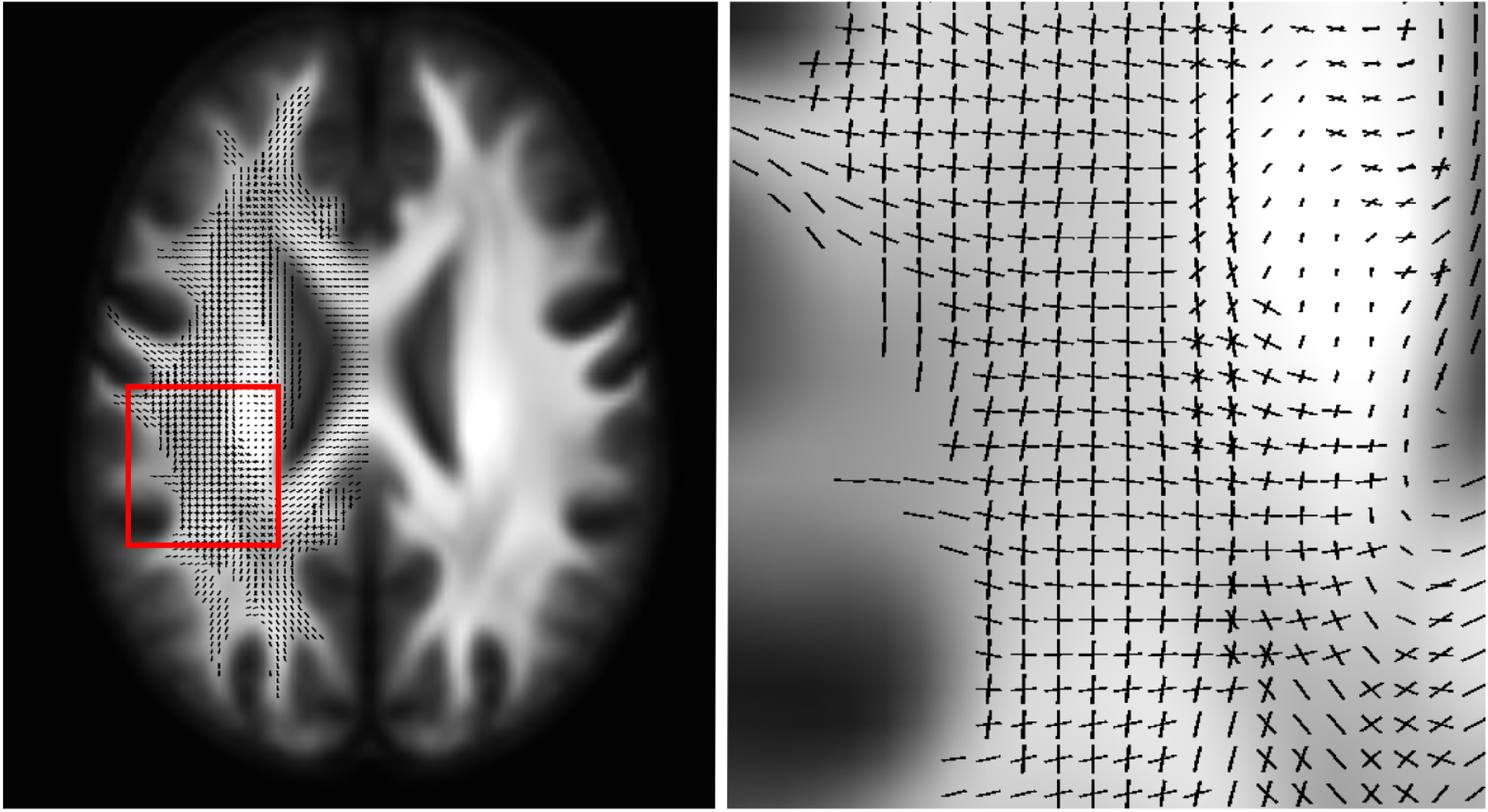
The final fixel analysis mask used for all statistical analyses in this work. It includes only fixels in a single hemisphere, since the laterality indices are computed from a pair of corresponding (left and right hemisphere) fixels in the brain. Here, this is visualised in the right hemisphere of the template (radiological convention). It is furthermore restricted to fixels for which correspondence with at least 90% of all participant hemisphere data could be established.

#### Fixel-wise statistical analyses

Non-parametric permutation testing (with 5000 permutations) was performed using connectivity-based fixel enhancement (Raffelt et al., 2015) within MRtrix3 (Tournier et al., 2019). Family-wise error (FWE) corrections were applied to control for false positives for all tests that were performed.

First, we separately tested the RLD group, the LLD group, as well as both groups combined (labelled “ALL”) for significant fixel-wise rightwards lateralisation (LI > 0) or leftwards lateralisation (LI < 0), and this also separately for each of the FD, FC and FDC metrics. This was achieved with one-sample t-tests (using random sign-flipping).

Second, a cross-sectional comparison between RLD and LLD was performed to examine *relative* group differences in lateralisation by comparing LI (of FD, FC and FDC) between both groups using two-sample t-tests. Note that a significant outcome of such a test can result from a range of underlying scenarios. For example, the scenario might be one of opposite directions of lateralisation between the groups. But it might also be a case of one group showing little to no lateralisation whereas the other one does. It could even result from both groups presenting with lateralisation in the same direction, but with a significantly different degree. Considering the scope of possible scenarios, the outcome of this test cannot trivially be derived from those of the aforementioned individual (one-sample) tests.

## Results

### Fibre-specific asymmetries in white matter

Figure 2 shows the result for the one-sample t-tests of LI(FDC) for the “ALL” (combined) group, as an example of how fixel-wise results of these tests can be understood as well as how they will be visualised in this work. The image in panel A and the zoomed regions in panels B and C present the combined *significant (p < 0*.*05)* fixel-wise results for both rightwards lateralisation (LI(FDC) > 0, in *orange*) and leftwards lateralisation (LI(FDC) < 0, in *blue*). The significant fixels are a subset of the fixel analysis mask (shown in Figure 1). Note that the FBA approach allows for fixel-specific results, whereby fixels related to specific white matter bundles can individually present as significant or not, even in the same voxel. In our laterality analyses, we found several regions where different crossing fibre populations in the same region (of voxels) even presented with significant lateralisation in *opposite* directions; i.e. regions where *orange* and *blue* coloured fixels cross. The visualisation using fixels themselves (as shown in Figure 2, panel B) is harder to inspect visually for large regions or entire 3D volumes. To mitigate this, a visualisation of the same result using a cropped streamlines tractogram (derived from the population FOD template) is used, as shown in Figure 2, panel C. Note this is merely an alternative visualisation of the same result, which has the benefit of appearing visually denser.

**Figure 2:**
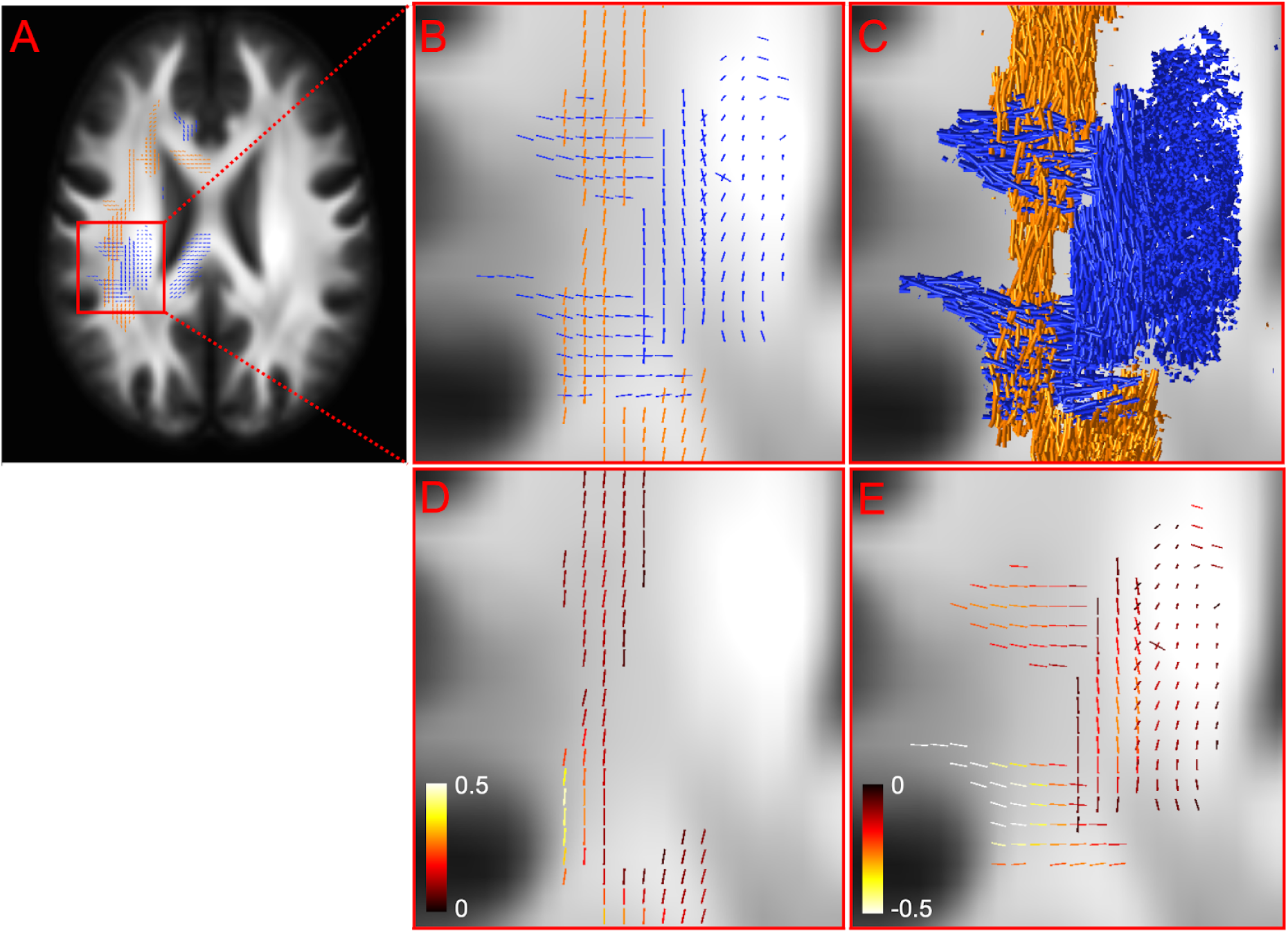
Fixel-wise results for the one-sample tests of LI(FDC) for the “ALL” (combined) group. Panels A-B: fixels with significant lateralisation are colour-coded. Orange indicates rightwards lateralisation (LI > 0) and blue indicates leftwards lateralisation (LI < 0). Panel C: a streamlines tractogram is cropped to display segments that correspond to significant fixels only. Panels D-E: effect sizes quantify the degree of lateralisation to the right and to the left hemisphere respectively.

In Figure 2, panels D and E, the effect size is shown for the significant fixels (rightwards and leftwards respectively). This represents the average LI(FDC) within the population (which is unbiased due to the additivity property of the log-ratio LI metric; see also Appendix A). For reference, an effect size of 0.5 (white on the hot colour scale used in Figure 2) represents an FDC_right_/FDC_left_ ratio of exp(0.5) = 1.65 (whereas red and yellow represent ratios of 1.18 and 1.40 respectively). The same holds symmetrically for the opposite direction of lateralisation.

Figure 3 provides an overview of the results of *all* one-sample t-tests for each group (ALL, LLD and RLD) and each FBA metric (FDC, FD and FC). The results are presented using the same style of visualisation employing a cropped streamlines tractogram, as in Figure 2, but in a full 3D glass brain volumetric visualisation. As before, all results are displayed within the right hemisphere. A detailed slice-wise overview of each separate result, as well as all effect sizes, is provided in the supplementary document.

**Figure 3:**
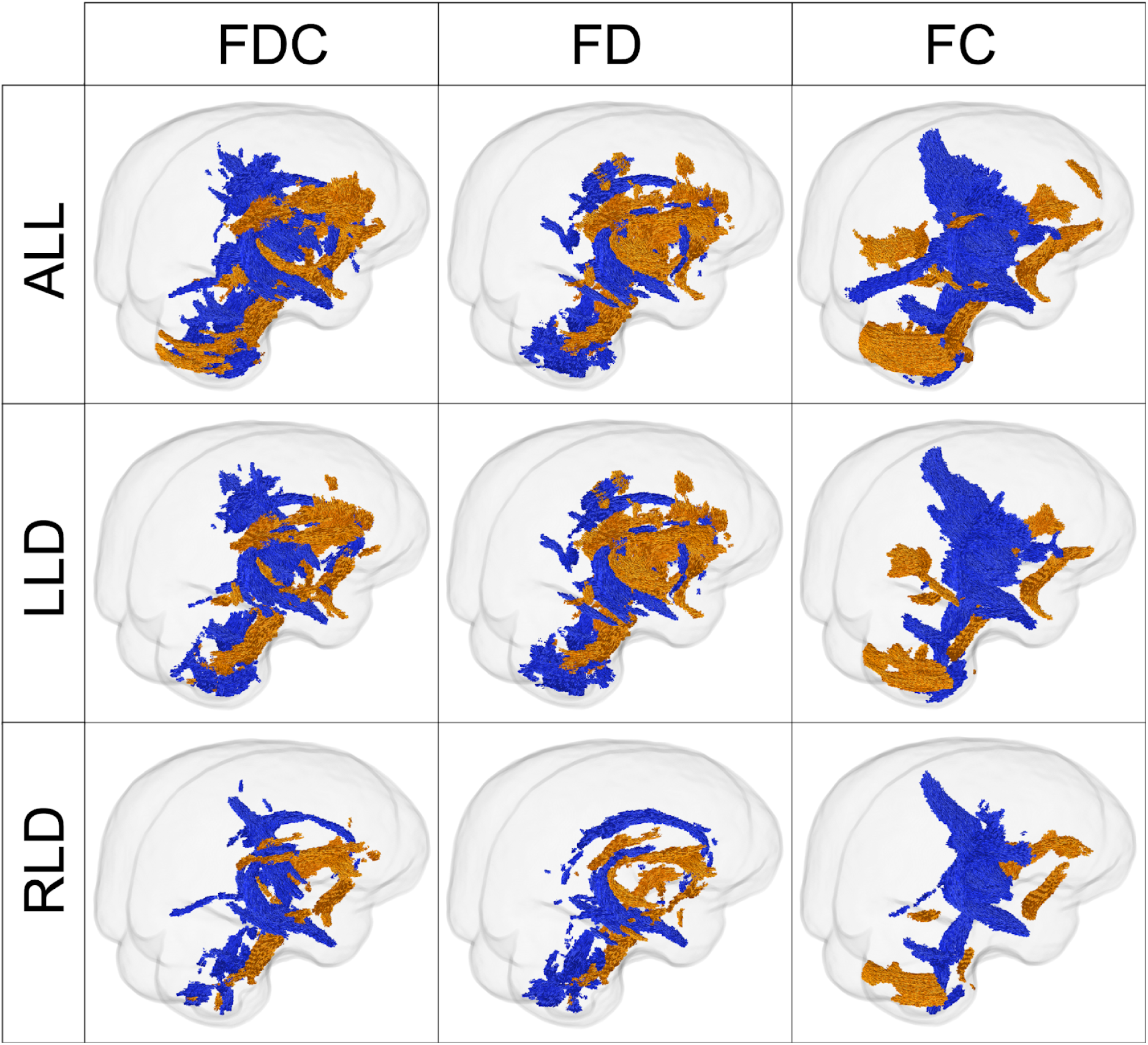
Visualisation of a cropped streamlines tractogram reflecting fixels with significant lateralisation in fibre density (FD), fibre-bundle cross-section (FC) and fibre density and cross-section (FDC) within a group of left language dominant (LLD) and right language dominant (RLD) participants and both groups combined (ALL). Streamlines coloured in orange are indicative of a rightwards lateralisation (LI > 0), whereas significant leftwards lateralisation is shown in blue (LI < 0).

The results for all groups and all metrics show large and extended parts of several white matter fibre bundles. Interestingly, a mix of rightwards and leftwards lateralisation of different structures is seen across the brain volume. Broadly, all groups show a very similar pattern of results, whereby the results appear slightly more extended in the LLD group as compared to the RLD group, as well as the ALL group compared to both others. This is in line with the relative numbers of participants in each group, whereby the analyses for larger sample sizes have more statistical power. The FDC results have some features in common with both FD as well as FC results, though appear to overall resemble the FD results slightly more.

The areas showing lateralisation of FDC to the right hemisphere include fixel clusters located in the anterior part of the arcuate fasciculus, the anterior limb of the internal capsule, the anterior and superior corona radiata, the body of the corpus callosum, the inferior-fronto-occipital fasciculus, the uncinate, the middle cerebral peduncle, the genu of the corpus callosum and part of the body of the corpus callosum.

Leftwards lateralisation of FDC was found in fixel clusters belonging to the long segment of the arcuate, inferior longitudinal fasciculus, cingulum (including the temporal part), fornix, external capsule, posterior parts of the body of the corpus callosum, inferior and superior cerebellar peduncles, posterior thalamic radiation, corticospinal tract, posterior parts of the superior corona radiata, the posterior limb of the internal capsule and the superior longitudinal fasciculus.

The results from the FD and FC analyses show both similar as well as additional lateralisation patterns. For FC, a rightwards lateralisation of the forceps major was found. Moreover, the leftwards lateralisation of FDC in the posterior limb of the internal capsule and the posterior part of the superior corona radiata is mainly supported by a lateralisation effect in FC. On the other hand, the leftwards lateralisation of FDC in the cingulum is only similar to a lateralisation effect in FD in the same structure.

### Relative group differences of fibre-specific asymmetries in white matter

The only significant difference between the LLD and RLD groups was found for LI(FC) in a cluster of fixels in the forceps minor, a white matter fibre bundle which connects the lateral and medial surfaces of the frontal lobes and crosses the midline via the genu of the corpus callosum (Figure 4, panel A). The difference is such that LI(FC) is significantly larger for the RLD group compared to the LLD group. Hence, this can be understood as the RLD group showing *relatively* more rightwards lateralisation compared to the LLD group.

**Figure 4:**
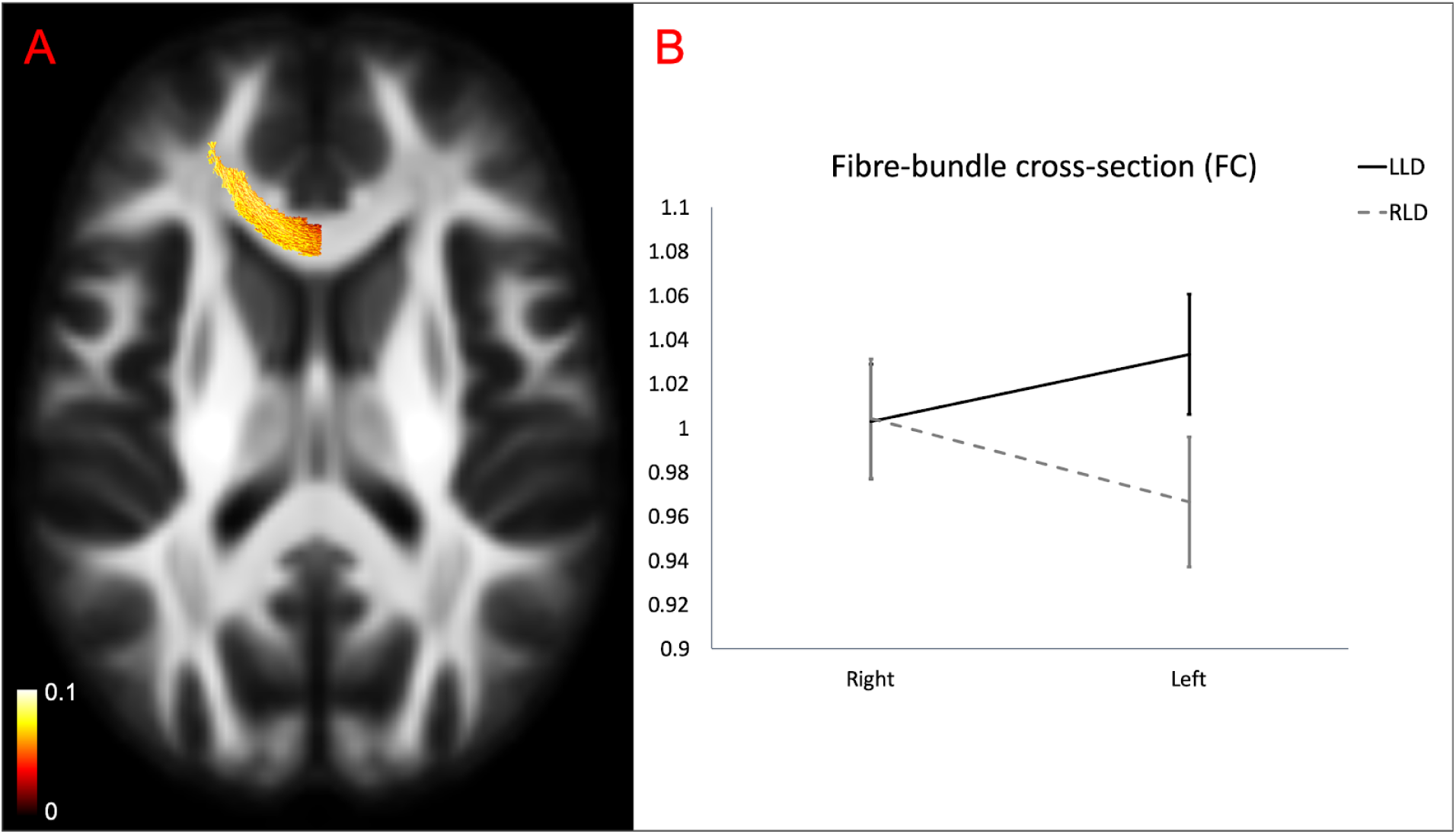
Panel A: Significant results for the two-sample test comparing the lateralisation index of fibre-bundle cross-section (LI(FC)) between the left language dominant (LLD) and the right language dominant (RLD) group. The group difference is such that LI(FC) is significantly larger for the RLD group compared to the LLD group. Fixels presenting with a significant group difference are visualised using a cropped streamlines tractogram and coloured by effect size. Positive values reflect a relatively stronger rightwards lateralisation in the RLD group compared to the LLD group. Panel B: Average FC values of the significant region of fixels for both groups and both hemispheres show on average a leftwards lateralisation of this region in the LLD group and a rightwards lateralisation in the RLD group. Error bars indicate the standard error.

As mentioned before, this effect can result from different kinds of underlying scenarios and we cannot trivially assume the lateralisation nor FC values themselves just from the outcome of this test alone. To better appreciate the full complexity of the present effect, we plotted the average (across the significant region of fixels) of FC values for both groups and both hemispheres (Figure 4, panel B). This reveals, on average, the nature of the effect: a rightwards lateralisation in the RLD group and a leftwards lateralisation in the LLD group. Moreover, both groups show similar FC values for the right side of the forceps minor, while the LLD group shows higher FC values on the left side compared to the RLD group.

Finally, note that the effect size (Figure 4, panel A) is expressed as the difference between the LI values for the RLD and LLD groups. Since this is a difference of two log-ratios, and due to the additivity property (Appendix A), it is itself a proper unbiased *relative* difference (and even a log-ratio: the logarithm of the ratio of the ratios).

## Discussion

This study examined white matter asymmetries, both on a macro- and microstructural level in a healthy sample of left-handed participants using a fixel-based analysis approach. Extensive areas of significant lateralisation were found in both directions, which were largely compatible with previous studies. The long segment of the arcuate fasciculus for example showed a leftwards lateralisation in both macro- and microstructure, as reported before in a voxel-based whole brain study (Ocklenburg et al., 2013), tract specific DTI studies (Bain et al., 2019; Catani et al., 2007; James et al., 2015), and in a recent FBA study (Arun et al., 2019). Similar to a study including left-handers by Vernooij et al. (2007) the corticospinal tract was lateralised to the left hemisphere. Other structures that showed asymmetries towards the left included part of the cingulum (analogous to (Arun et al., 2019; Slater et al., 2019)) and the inferior longitudinal fasciculus (comparable to (Arun et al., 2019; Thiebaut de Schotten, ffytche, et al., 2011)). With regard to the uncinate, a rightwards lateralisation was found, which agrees with previous findings (Arun et al., 2019; Highley et al., 2002; Park et al., 2004). Additionally, a rightwards lateralisation was observed for parts of the inferior fronto-occipital fasciculus and the anterior segment of the arcuate fasciculus.

From these results it is clear that white matter distribution, both at microscopic as well as macroscopic scales, is far from symmetric in the human brain. This widespread structural hemispheric asymmetry has important implications for clinical studies where patients’ images are often flipped across the midline to align the affected hemispheres, for example in studies with temporal lobe epilepsy or stroke. Without caution, this practice might result in false positive findings related to typical asymmetries such as those found in our current work. First of all, it is likely important to flip the same proportion of images of healthy control groups in order to match the distributions of structural asymmetries. Secondly, however, this will likely increase variability in all subject groups in such analyses, and thus decrease the power to detect subtle group differences. Finally, effects of lateralised pathologies might themselves of course also present differently depending on the direction of laterality. They might even interact differently with the existing lateralised structure of white matter in both hemispheres. Taking all of this into account, it might thus be advisable to even avoid the brain flipping strategy altogether in such studies, and rather analyse both patient groups separately. The latter approach was for example taken by Egorova et al. (2020), who compared left and right side infarcted stroke groups *separately* to controls, to allow for greater specificity in results.

Studies examining the relationship between left-right asymmetries and language functioning to date, have almost exclusively included left language dominant (LLD) participants (Ocklenburg et al., 2016). They have reported significant correlations between brain activation during language tasks and structural asymmetries based on diffusion MRI data, and concluded that these anatomical asymmetries may reflect or even cause functional language laterality. However, by not including right language dominant (RLD) participants, this hypothesized structure-function relationship has remained speculative. To further uncover the relationship between functional and structural language lateralisation, our present study included a relatively large sample of participants with RLD. Surprisingly few differences were found when comparing whole brain lateralisation patterns of white matter between the LLD and RLD groups. Moreover, no between-group differences of structural asymmetry were found in the arcuate fasciculus. This is in line with the study by Vernooij et al. (2007) that showed a leftwards lateralisation of the arcuate regardless of the direction of functional hemispheric dominance for language. This suggests that the lateralisation of language functioning and the arcuate fasciculus are largely driven by independent biases. Finally, a significant between-group difference in laterality of fibre-bundle cross-section (FC) was found in the forceps minor. Interestingly, FC of the right hemisphere in this region was, on average, actually very similar in both groups, while a difference was observed in the left hemisphere, leading to opposite directions of laterality (i.e. leftwards in the LLD and rightwards in the RLD group). The forceps minor, also known as the anterior forceps, is a white matter fibre bundle which crosses the midline via the genu of the corpus callosum and connects the lateral and medial surfaces of the frontal lobes. Contralateral connections travel via the corpus callosum with the forceps minor to end in Brodmann areas 9 and 10. These areas are involved in executive control and working memory (area 9) and in abstract cognitive functioning, episodic and working memory (area 10) (Baker et al., 2018). A large DTI twin study found no genetic influence for its leftward laterality, but shared environmental factors account for about 10% of its variance in asymmetry (Jahanshad et al., 2010). The exact function of the forceps minor itself remains elusive. Fractional anisotropy (derived from DTI) shows a positive correlation with expressive language and cognitive control abilities in children in this tract (Farah et al., 2020). Other DTI-based findings in adults suggest a relation with indices of general and crystallised intelligence (Cox et al., 2019; Góngora et al., 2020). In agreement with these findings, recent clinical research also reports that white matter hyperintensities in the forceps minor predict global cognitive impairment (Biesbroek et al., 2020). The establishment of a complete and formal account of the relation between forceps minor asymmetry and language dominance requires further investigation.

Several diffusion MRI studies have previously reported on structural white matter asymmetries using metrics derived from the DTI model (Güntürkün & Ocklenburg, 2017; Ocklenburg et al., 2016). By using the FBA approach in our current study, important limitations inherent to the DTI model were resolved, leading to more reliable and informative results. FBA allows for statistical analysis of fibre-specific measures (Raffelt et al., 2017). In this work, we combined the FBA framework with a bespoke laterality index, computed for the 3 traditional FBA metrics (FD, FC and FDC). As a result, we were able to determine lateralisation of multiple fibre populations within a voxel individually. In our work, this specificity of the FBA framework shows in several regions of the results, including for example fixels within the arcuate showing opposite asymmetry (Figure 2). The long segment was lateralised to the left side of the brain, whereas fixels belonging to the anterior part and crossing the long segment showed a rightwards lateralisation. Voxel-averaged DTI measures might have failed to detect and explain the lateralisation pattern within these and similar regions containing multiple crossing fibre populations, underscoring the added value of using FBA for laterality research of white matter. Thanks to the FBA metrics it was furthermore possible to distinguish between macroscopic and microscopic asymmetries of white matter properties. For example, the cingulum showed a clear leftwards lateralisation of fibre density, sensitised to axonal volume at a microscopic scale. This asymmetry was not present for the fibre-bundle cross-section at the macroscopic scale. Such a result is for example compatible with the leftward lateralisation of FA for the cingulum, which was found by previous studies (Gong et al., 2005; Park et al., 2004), while Thiebaut de Schotten, ffytche et al. (2011) also failed to find volume differences in the left and right cingulum in most subjects of their study. Finally, FBA is a whole brain data-driven analysis approach allowing to detect structural asymmetries in subparts of white matter tracts without any a priori hypotheses. This is in contrast to several studies examining white matter lateralisation via delineation of tracts of interest and extraction of region-wise averaged metrics, which lack regionally varying specificity to detect subtle effects.

The FBA statistical framework has been used in many different studies over the past few years. Whole brain white matter asymmetry in a healthy adult sample has been examined using the FBA approach by Arun et al. (2019). While they employed one-sample testing of the *absolute* difference between left and right hemisphere FBA measures, we opted for a bespoke *relative* laterality index to more directly quantify asymmetries specifically. This represents an approach commonly used in the broader field of lateralisation research (Seghier, 2008). It also allowed us to compare the relative degree of laterality directly and in an intuitive manner *between* different populations. However, rather than directly adopting the most common definition of a relative laterality index—the difference divided by the sum (Seghier, 2008)—we defined our laterality index as a log-ratio. While the traditional laterality index is symmetric, it is also bounded (between values of −1 and 1), and therefore lacks the additivity property. The log-ratio, on the other hand, is the only measure of relative difference that is not only symmetric, but also normed and additive (Tornqvist et al., 1985). Despite its definition potentially appearing quite different at first sight, a direct and intuitive relationship exists between both indices (Appendix A). Our log-ratio laterality index can be calculated directly on a fixel-specific level, though particular care has to be taken in implementing strategies to deal with “absent” fibre densities (however, this is not unlike the strategy to assign these a value of zero in traditional FBA studies).

One of the limitations of the present study is the inclusion of left-handed participants only. To increase the probability of finding right language dominant participants, we focused on left-handers because RLD is rare and occurs more often in this population (Carey & Johnstone, 2014). As a result of this approach, an unprecedented sample of RLD participants was recruited. The question however remains whether structural asymmetries are comparable in right-handed LLD and RLD.

## Conclusion

We found that white matter asymmetries are ubiquitous throughout the brain, which holds important consequences for clinical research. Fixel-based analyses present a promising avenue to study the asymmetry of white matter using a fibre-specific laterality index defined for micro- and macrostructural metrics. The direction of language lateralisation appeared to be unrelated to the structural asymmetry of white matter tracts known to be typically involved in language. Finally, a significant difference in relative lateralisation was found for part of the forceps minor when comparing participants with left and right language dominance. The exact relationship between forceps minor asymmetry and language dominance could be an interesting subject of future studies.

## Appendix A

In lateralisation research, traditionally a measure of *relative* difference is used to represent the direction and degree of laterality independently of the absolute magnitude of relevant measurements (e.g. cortical volume and others). This has the benefit of intuitively quantifying asymmetries specifically, and allows for direct comparison of these between different structures or populations. The “traditional” laterality index (LI_trad_) used for this purpose is calculated as the difference divided by the sum, sometimes multiplied by a constant (Seghier, 2008). In the most commonly appearing variant, this constant is simply set to 1, resulting in

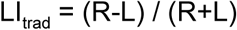

with R and L here representing the relevant right and left side measurements. This is a *symmetric* version of the typical definition of a relative difference, which would otherwise divide by just one of both values (i.e. R *or* L), resulting in an asymmetric definition which is obviously undesirable for expressing laterality in an intuitive and unbiased manner. Interestingly, the above LI_trad_ is *not* properly normed; it *would* be if the sum in the denominator were a *mean* instead, i.e. by introducing a factor 2 (Tornqvist et al., 1985). This would render it more intuitive in particular for smaller values, which can then be thought of directly as a “percentage” difference, yet without sacrificing the valuable symmetry property.

However, in this work we defined our laterality index (LI) as a log-ratio instead, i.e.,

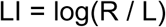

where “log” refers to the natural logarithm. This measure of relative difference is also symmetric, as well as properly normed (Tornqvist et al., 1985). However, it’s also *additive*: log-ratios can be added directly to calculate unbiased total changes (i.e. sums of changes), which themselves also represent a proper log-ratio. The aforementioned LI_trad_ *lacks* additivity: this can be easily seen by the fact that it is bounded (between −1 and 1).

While the definitions of LI_trad_ and LI may appear quite different at first sight, they do feature a direct relationship due to the fact that they are both measures of *relative* difference. All relative difference measures can ultimately be written in function of the ratio between both values (e.g. R and L), independently of the magnitude of those values themselves. This allows us to express LI_trad_ directly in function of LI:

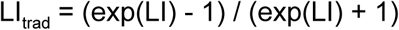

To aid intuition, this relationship is plotted in Figure 5. This reveals that, for small magnitudes, both indices are almost linearly related (via the aforementioned factor 2). While our *average* effect sizes are only in an (LI) range of −0.5 to 0.5, note that the original subject-wise values will vary more. Hence, the additivity property is key in calculating these very averages in an unbiased manner. Similarly, relative comparisons of laterality between populations rely on computing differences, which can also only be done in an unbiased way when the additivity property is satisfied.

More details are provided in Tornqvist et al. (1985). A comparison of the values of LI and other relative measures, for the range of (average) effect sizes in this work, is provided in Table 1.

**Figure 5:**
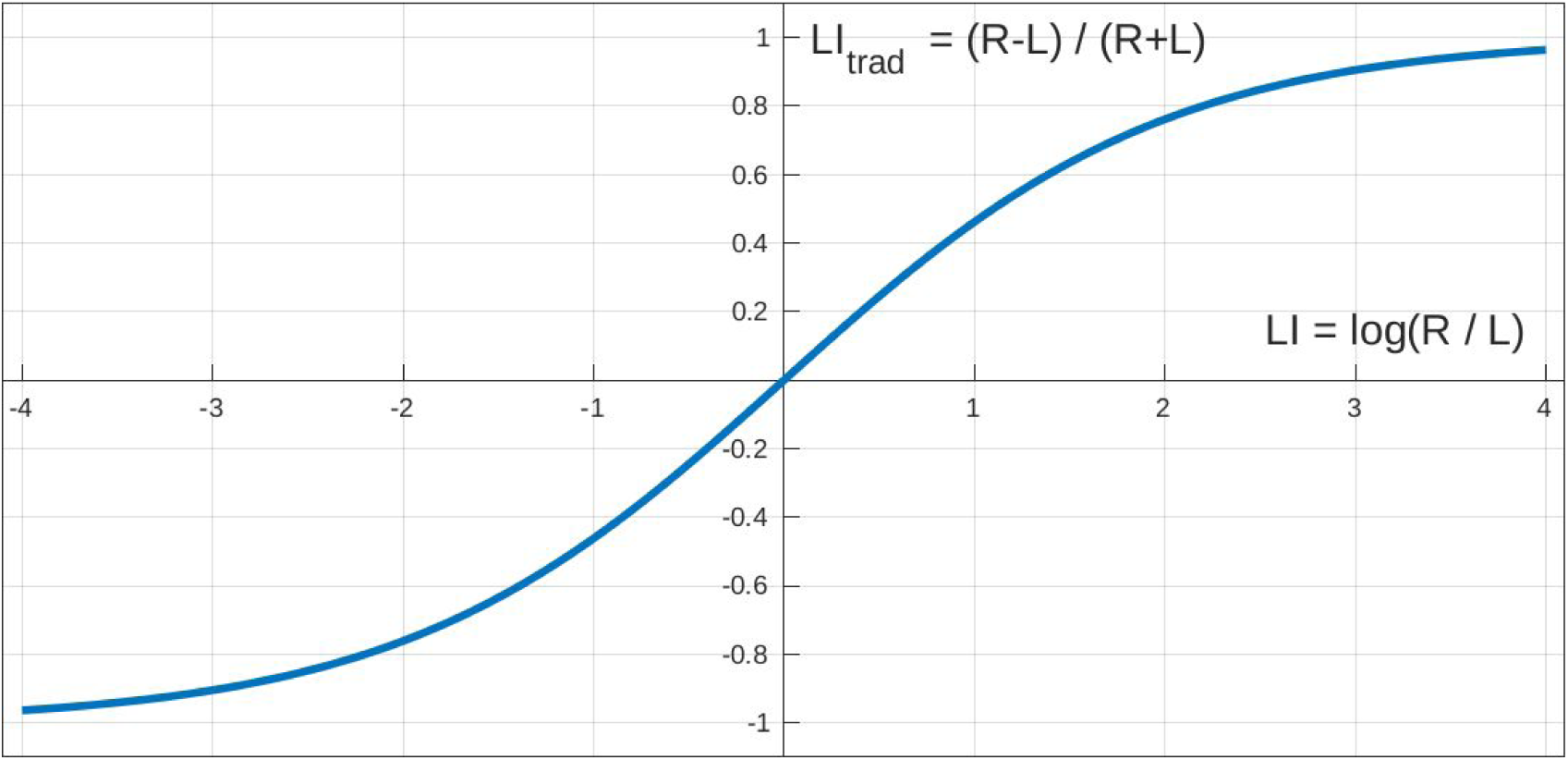
The traditional laterality index (difference divided by sum) in function of our laterality index (a log-ratio). Because both are measures of relative difference, their values can be related to each other independently of the absolute values of R and L.

**Table 1:** Corresponding values of other measures of relative difference, for the range of values of our log-ratio LI as used to describe effect sizes in the figures in this work and the supplementary document (from 0.5 down to −0.5).

| $LI = \log(R / L)$ | $R / L$ | $(R-L) / L$ | $(R-L) / R$ | $(R-L) / ((R+L) / 2)$ | $LI_{\text{trad}} = (R-L) / (R+L)$ |
| --- | --- | --- | --- | --- | --- |
| 0.5 | 1.649 | 0.649 | 0.393 | 0.490 | 0.245 |
| 0.4 | 1.492 | 0.492 | 0.330 | 0.395 | 0.197 |
| 0.3 | 1.350 | 0.350 | 0.259 | 0.298 | 0.149 |
| 0.2 | 1.221 | 0.221 | 0.181 | 0.199 | 0.100 |
| 0.1 | 1.105 | 0.105 | 0.095 | 0.100 | 0.050 |
| 0.0 | 1.000 | 0.000 | 0.000 | 0.000 | 0.000 |
| -0.1 | 0.905 | -0.095 | -0.105 | -0.100 | -0.050 |
| -0.2 | 0.819 | -0.181 | -0.221 | -0.199 | -0.100 |
| -0.3 | 0.741 | -0.259 | -0.350 | -0.298 | -0.149 |
| -0.4 | 0.670 | -0.330 | -0.492 | -0.395 | -0.197 |
| -0.5 | 0.607 | -0.393 | -0.649 | -0.490 | -0.245 |

