## Supplementary document for "Fibre-specific laterality of white matter in left and right language dominant people"

### Supplemental Materials

A detailed slice-wise overview of all results is presented here. To enable the visualisation of the significant fixels, streamlines from the population FOD template tractogram were cropped to include streamline points that correspond to significant fixels (FWE-corrected p-value <0.05).

For each one-sample t-test the following is visualised:

- combined significant results for both rightward lateralisation (LI > 0, in *orange*) and leftward lateralisation (LI < 0, in *blue*).
- effect size of the significant results defined as average LI within the population.

ALL = all participants together (RLD and LLD)
RLD = right language dominant participants
LLD = left language dominant participants

FDC = fibre density and cross-section
FD = fibre density
FC = fibre-bundle cross-section

**INDEX**


ALL - FDC - direction of lateralisation 2
ALL - FDC - effect size - both directions 3
ALL - FDC - effect size - right lateralisation 4
ALL - FDC - effect size - left lateralisation 5
ALL - FD - direction of lateralisation 6
ALL - FD - effect size - both directions 7
ALL - FD - effect size - right lateralisation 8
ALL - FD - effect size - left lateralisation 9
ALL - FC - direction of lateralisation 10
ALL - FC - effect size - both directions 11
ALL - FC - effect size - right lateralisation 12
ALL - FC - effect size - left lateralisation 13
LLD - FDC - direction of lateralisation 14
LLD - FDC - effect size - both directions 15
LLD - FDC - effect size - right lateralisation 16
LLD - FDC - effect size - left lateralisation 17
LLD - FD - direction of lateralisation 18
LLD - FD - effect size - both directions 19
LLD - FD - effect size - right lateralisation 20
LLD - FD - effect size - left lateralisation 21
LLD - FC - direction of lateralisation 22
LLD - FC - effect size - both directions 23
LLD - FC - effect size - right lateralisation 24
LLD - FC - effect size - left lateralisation 25
RLD - FDC - direction of lateralisation 26
RLD - FDC - effect size - both directions 27
RLD - FDC - effect size - right lateralisation 28
RLD - FDC - effect size - left lateralisation 29
RLD - FD - direction of lateralisation 30
RLD - FD - effect size - both directions 31
RLD - FD - effect size - right lateralisation 32
RLD - FD - effect size - left lateralisation 33
RLD - FC - direction of lateralisation 34
RLD - FC - effect size - both directions 35
RLD - FC - effect size - right lateralisation 36
RLD - FC - effect size - left lateralisation 37

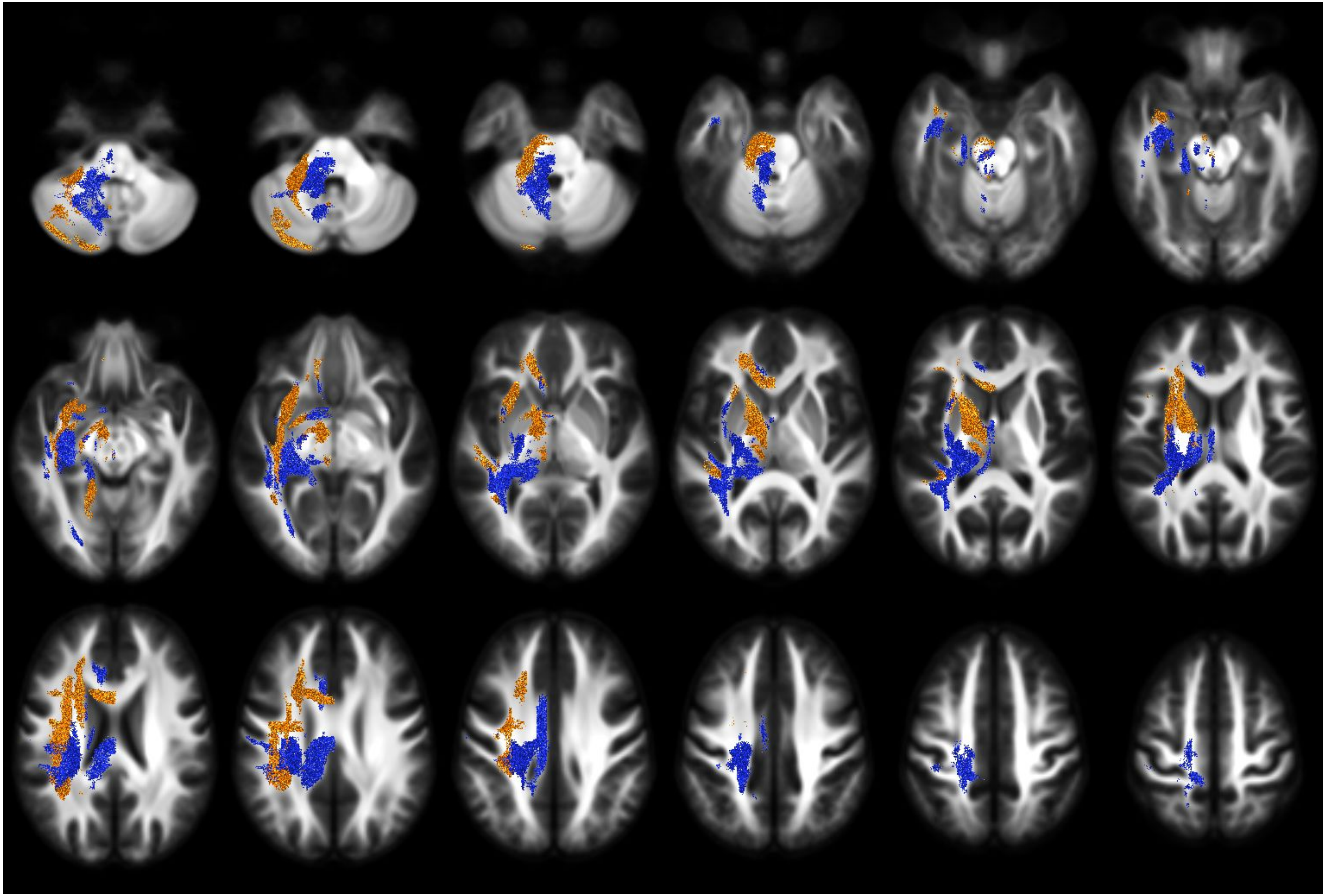

ALL - FDC - direction of lateralisation

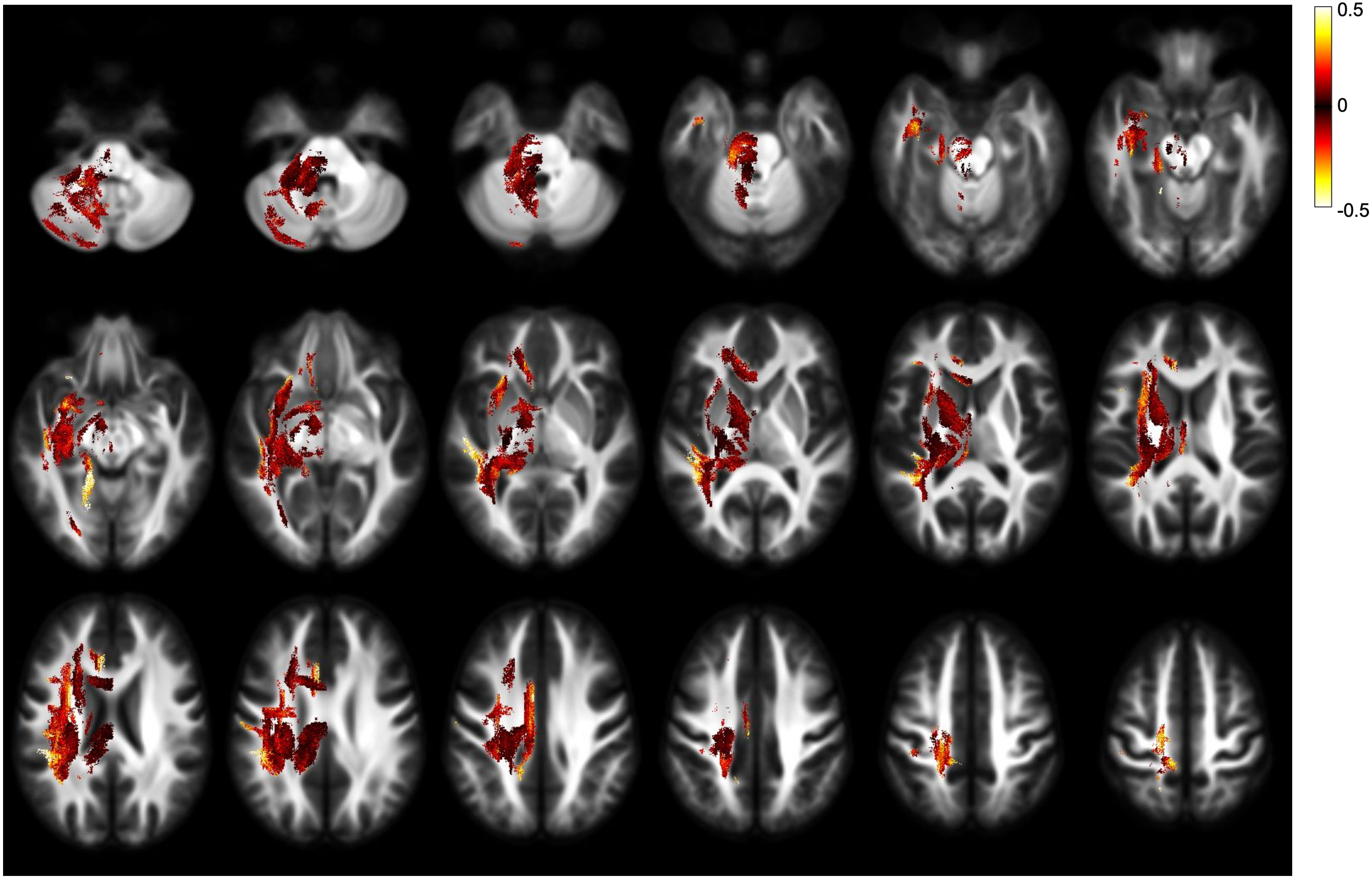


ALL - FDC - effect size - both directions

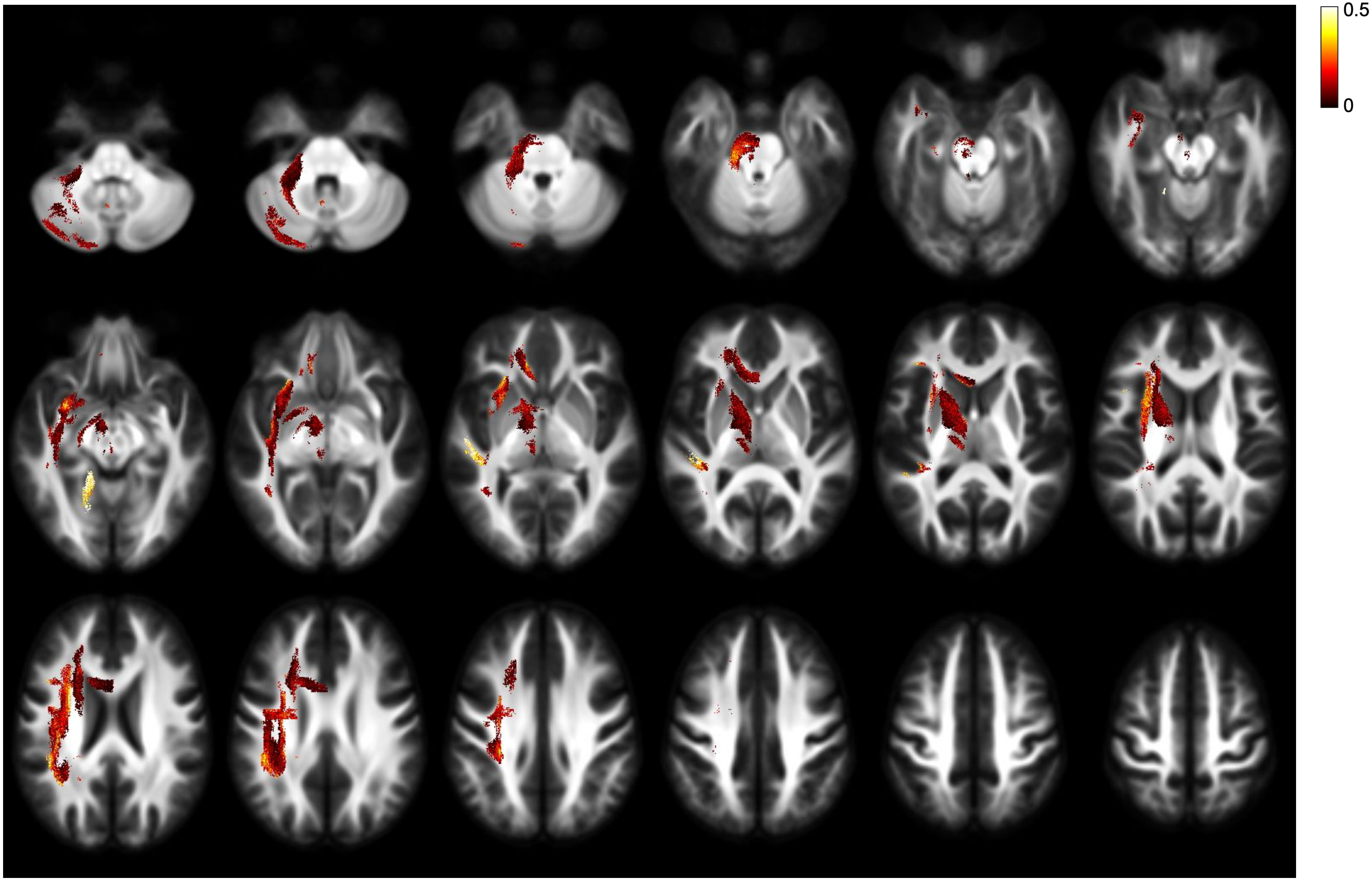


ALL - FDC - effect size - right lateralisation

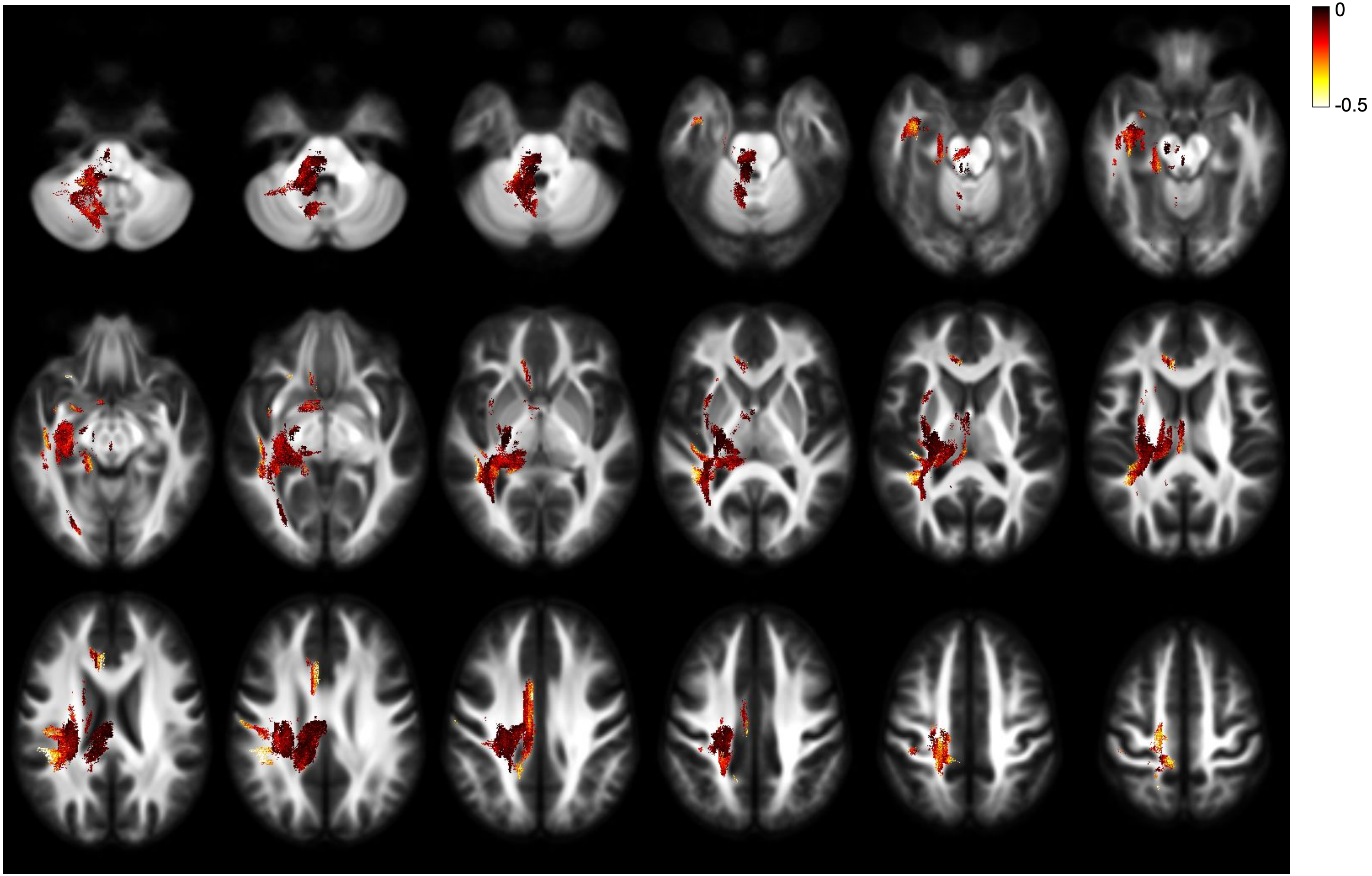


ALL - FDC - effect size - left lateralisation

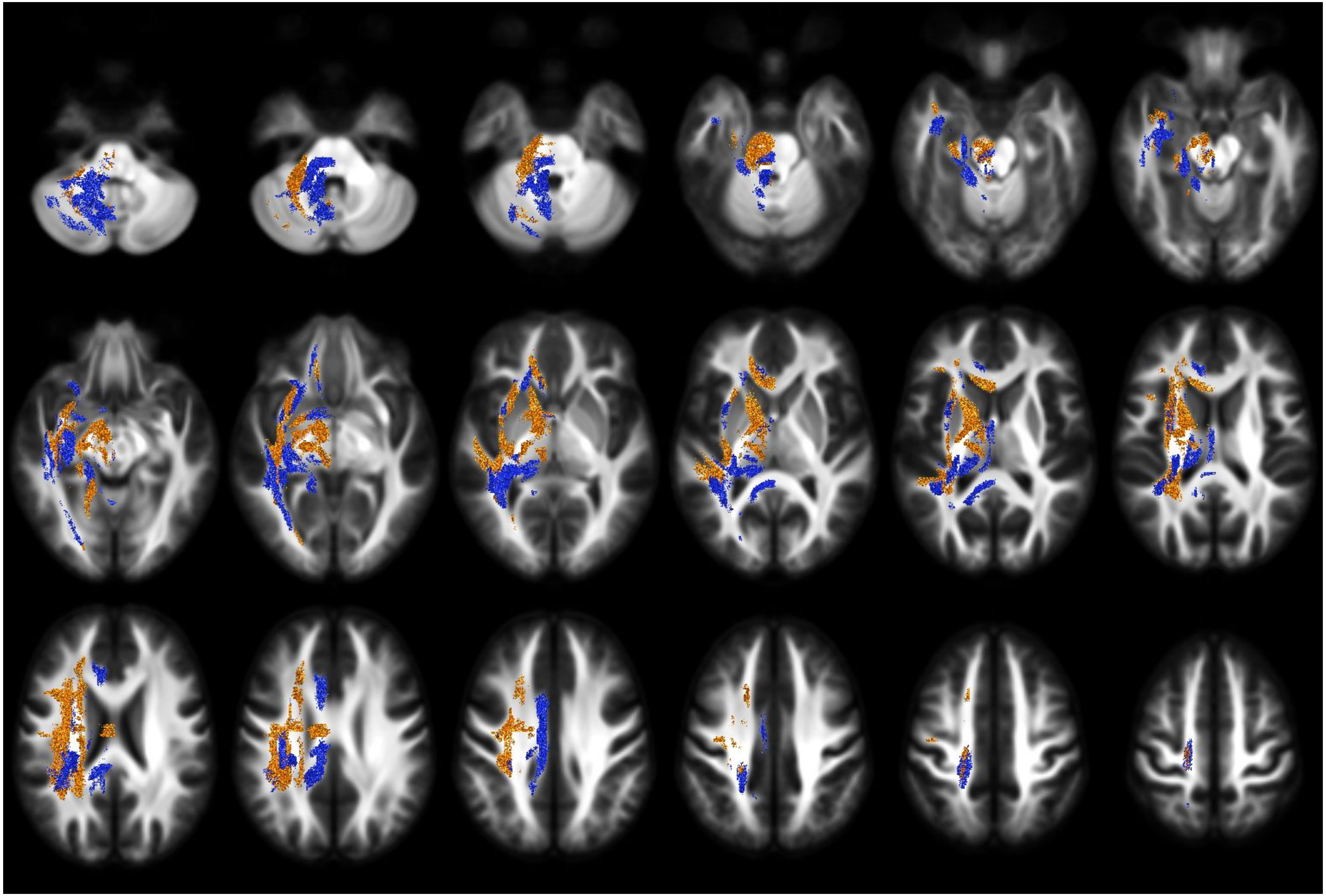

ALL - FD - direction of lateralisation

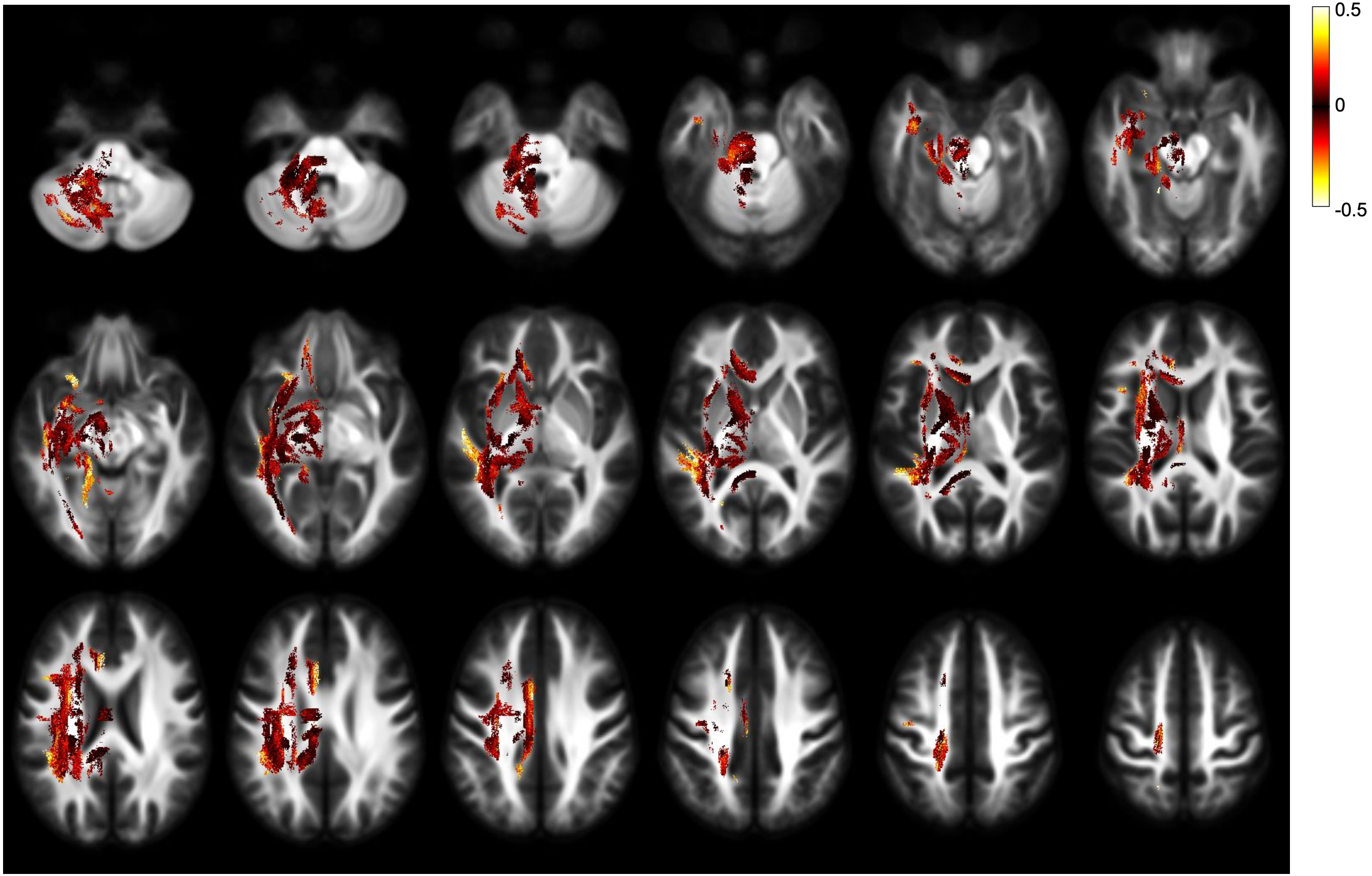


ALL - FD - effect size - both directions

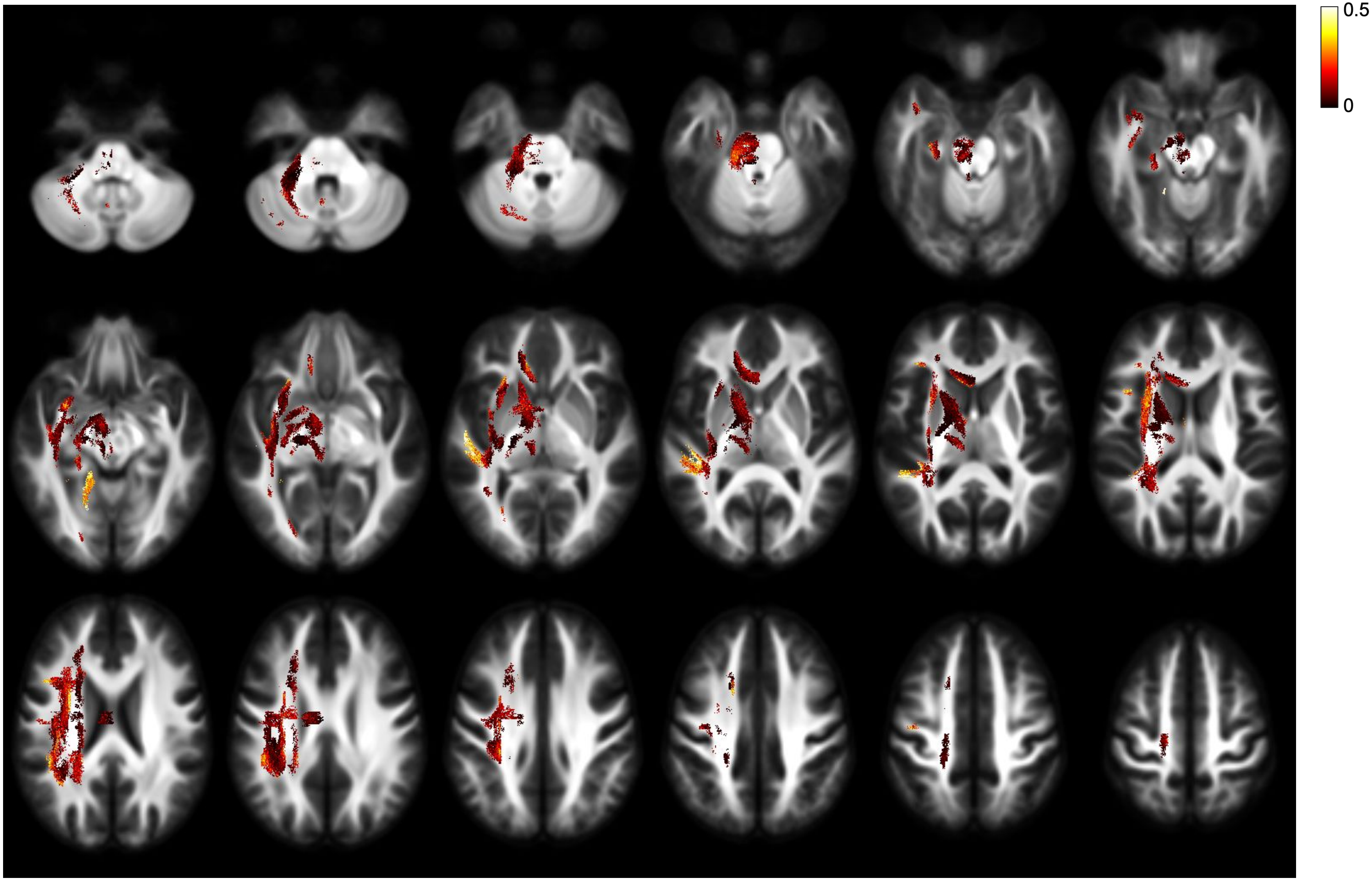


ALL - FD - effect size - right lateralisation

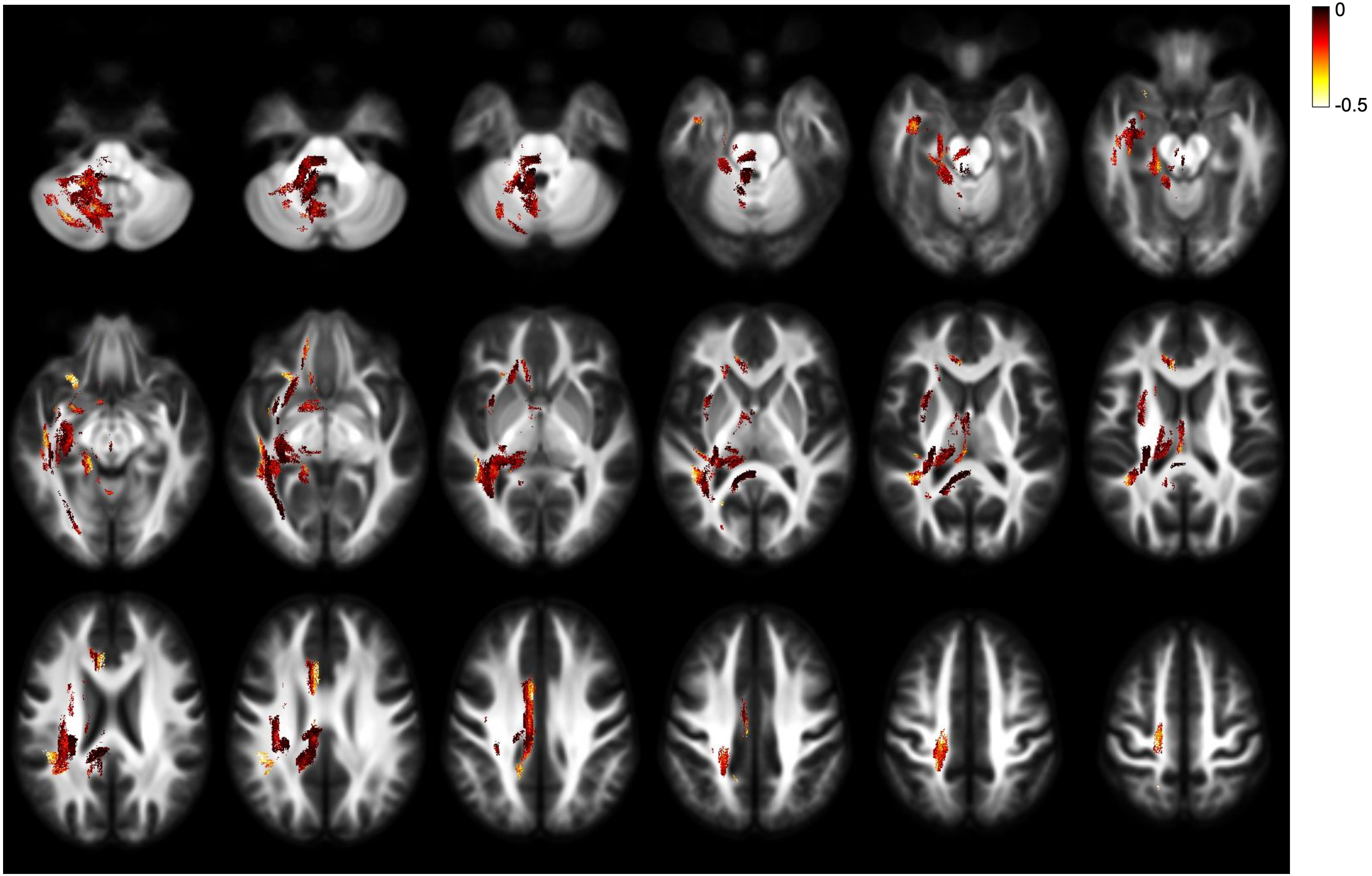


ALL - FD - effect size - left lateralisation

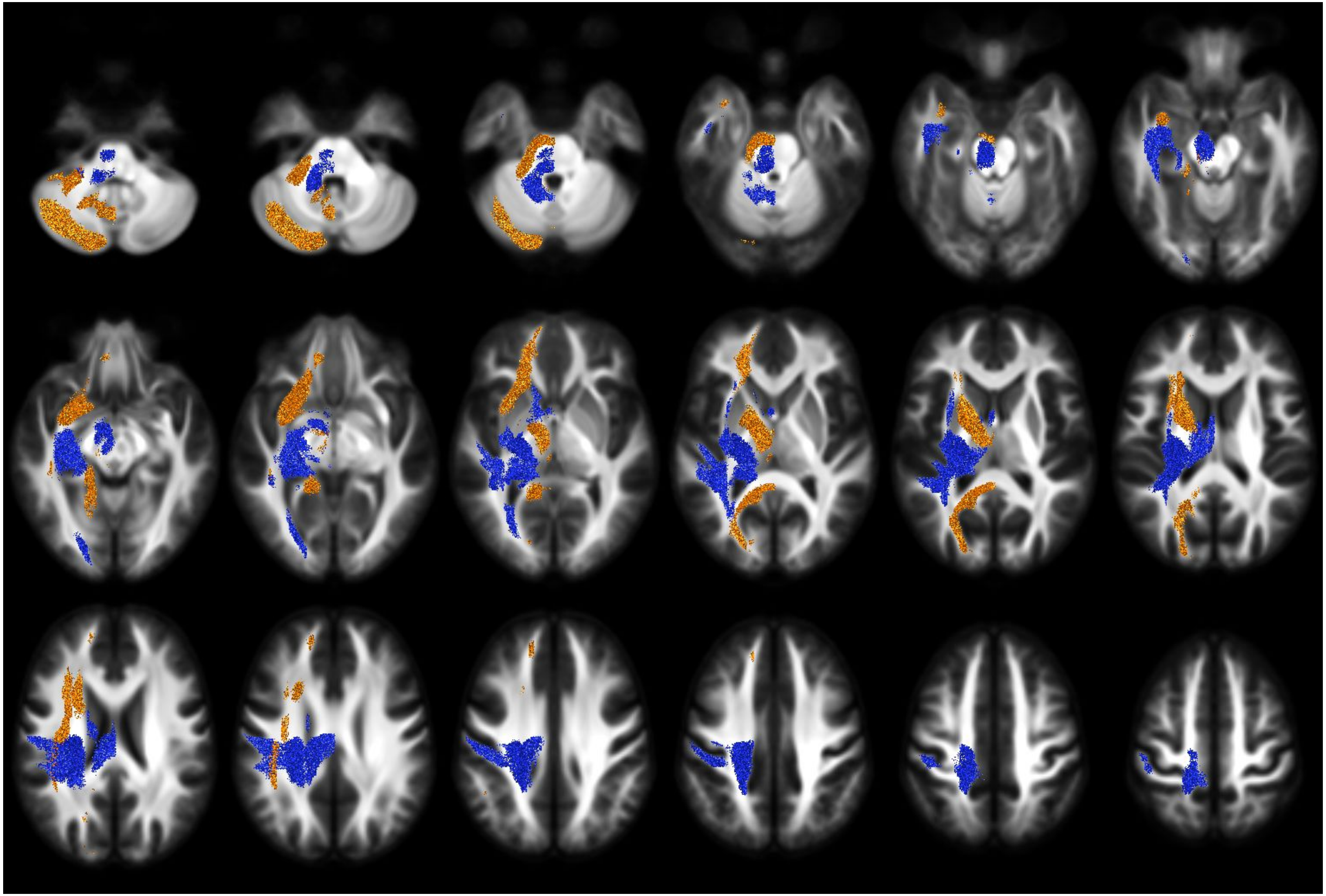

ALL - FC - direction of lateralisation

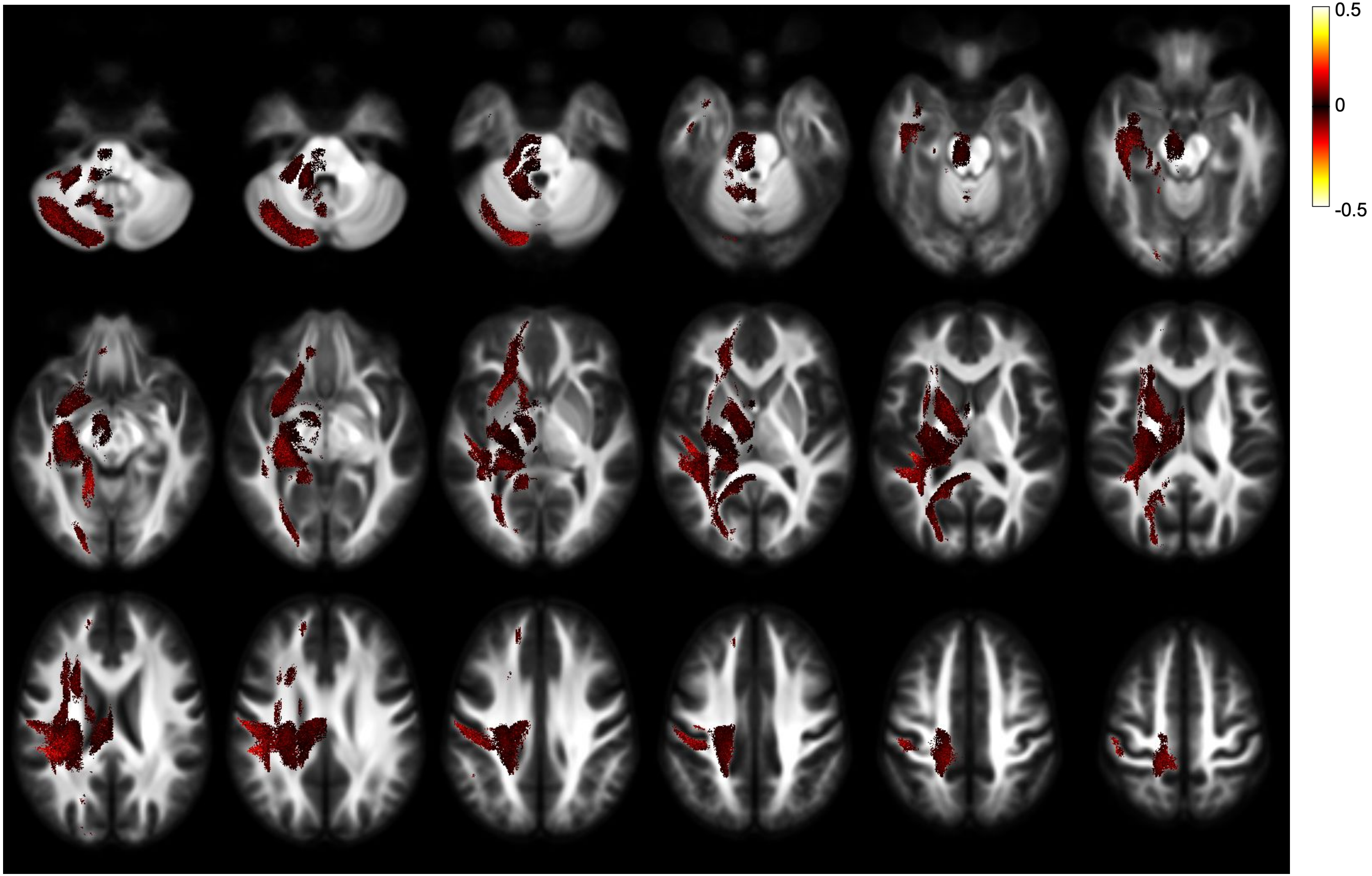


ALL - FC - effect size - both directions

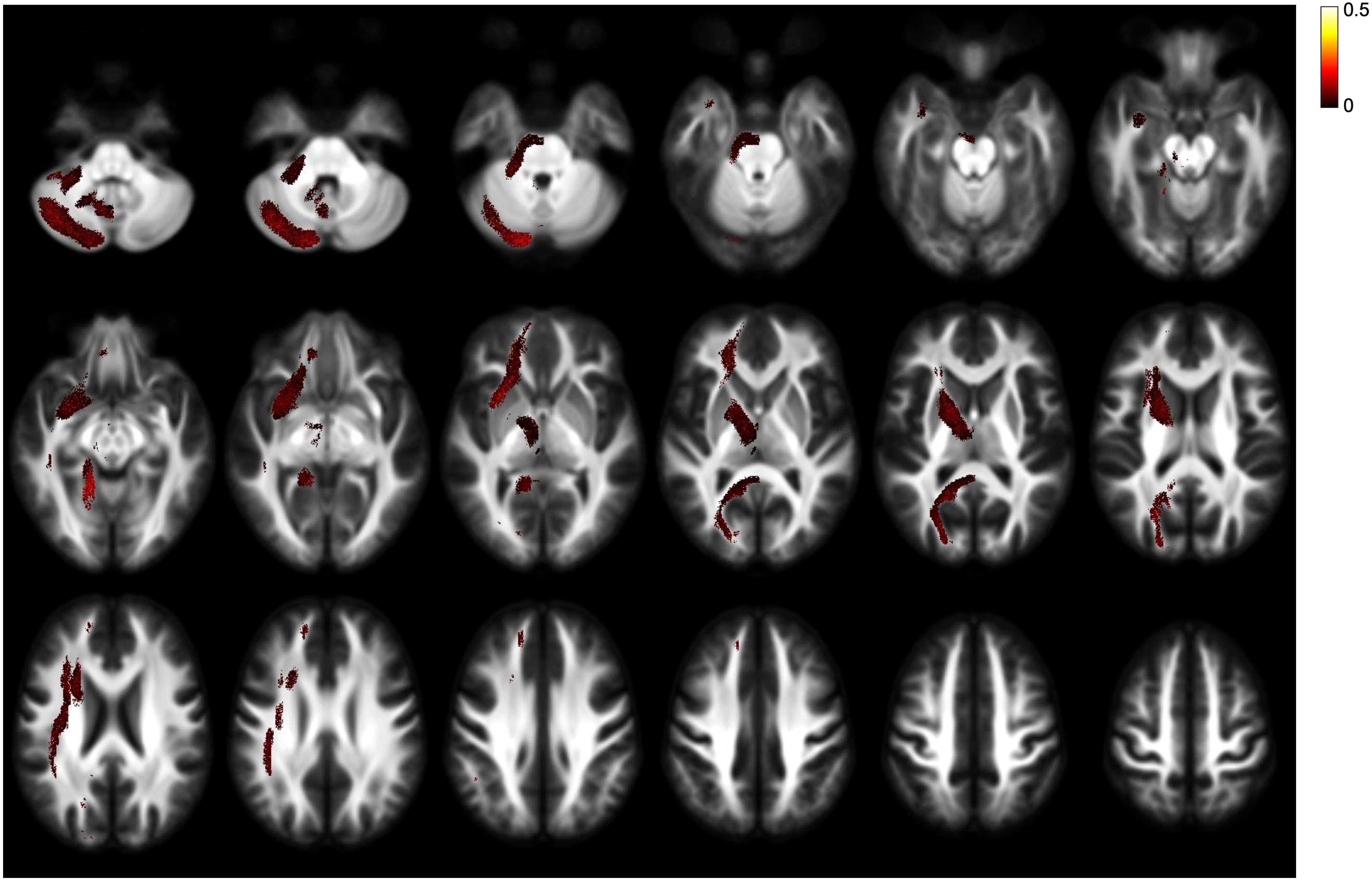


ALL - FC - effect size - right lateralisation

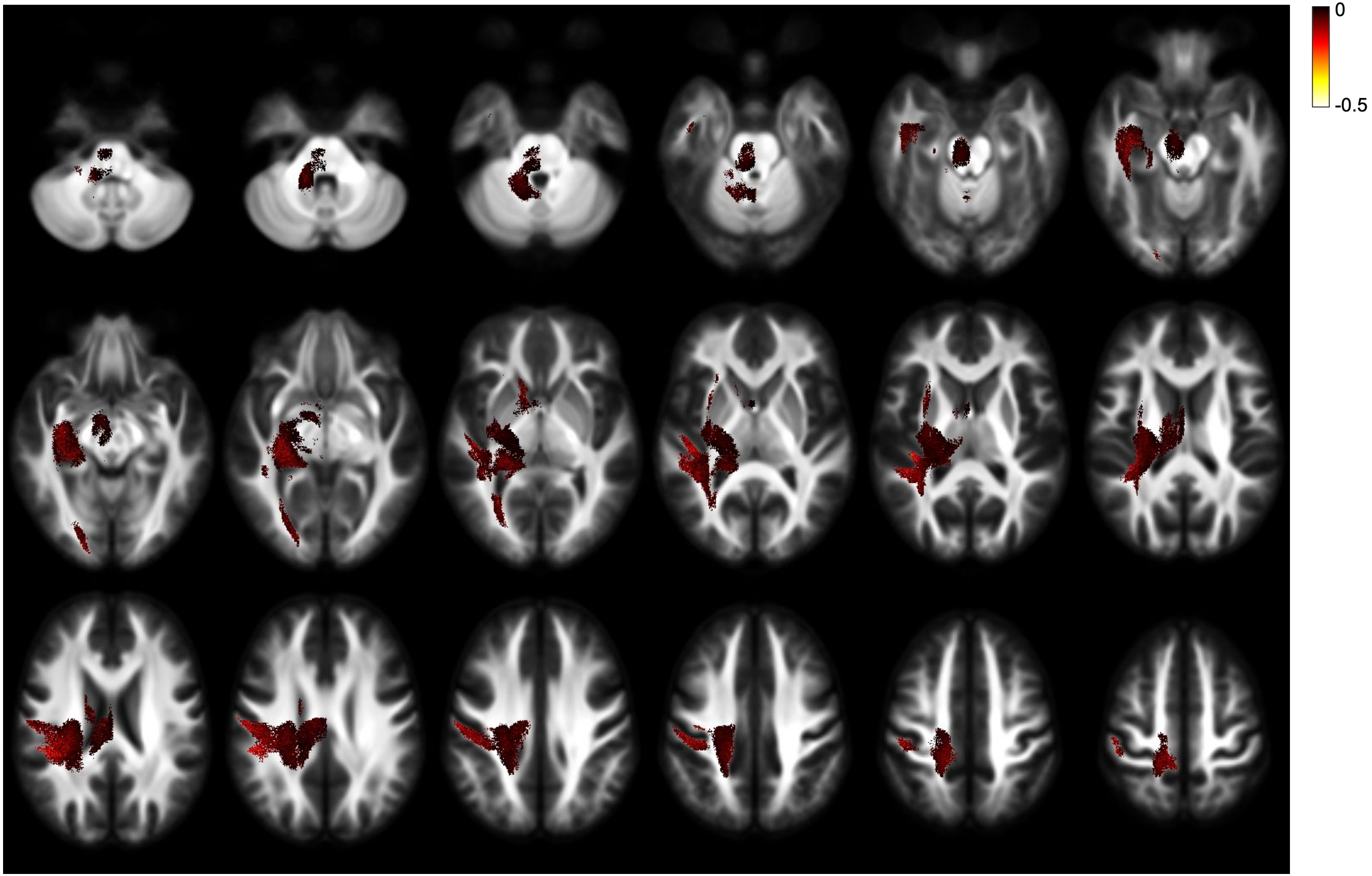


ALL - FC - effect size - left lateralisation

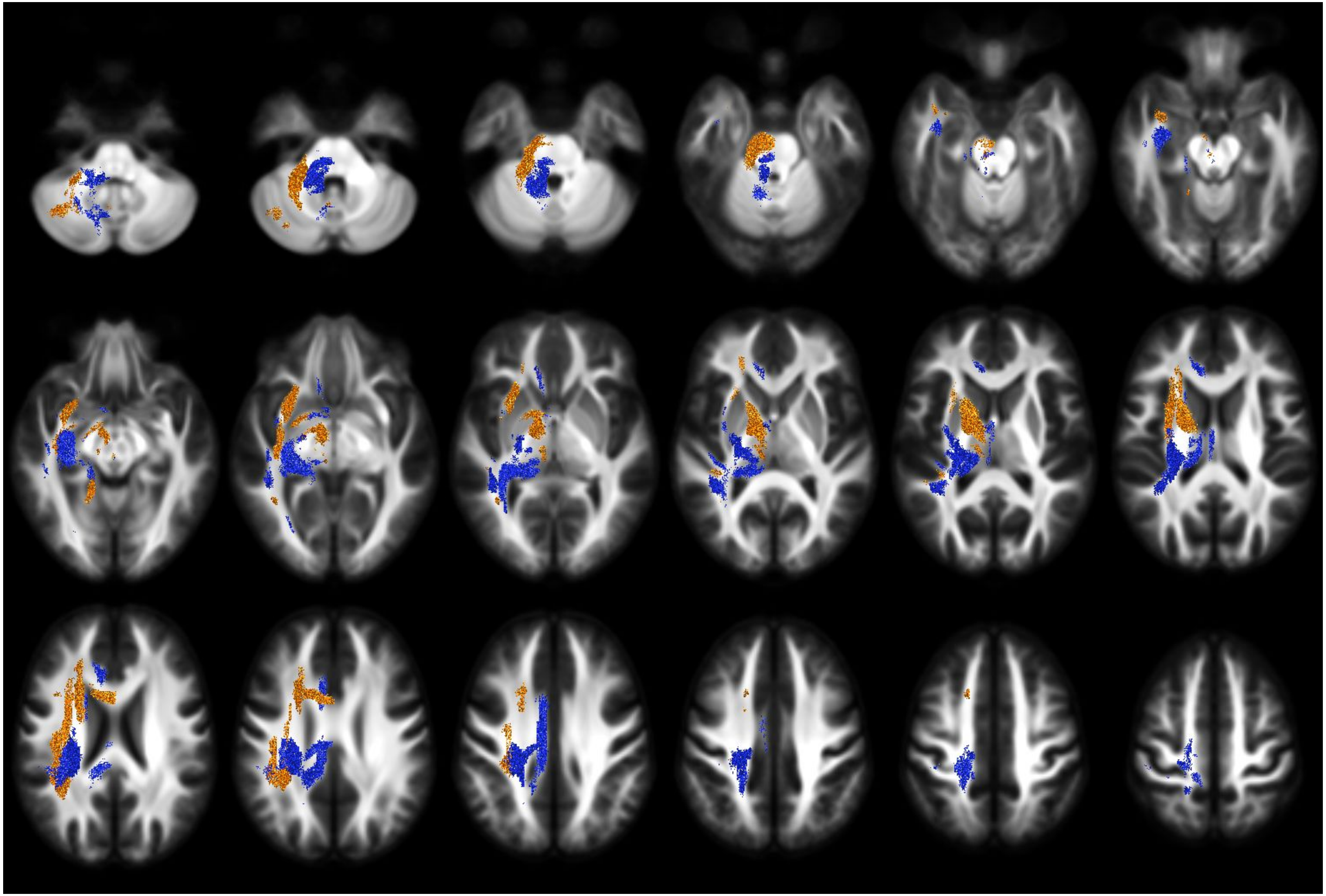

LLD - FDC - direction of lateralisation

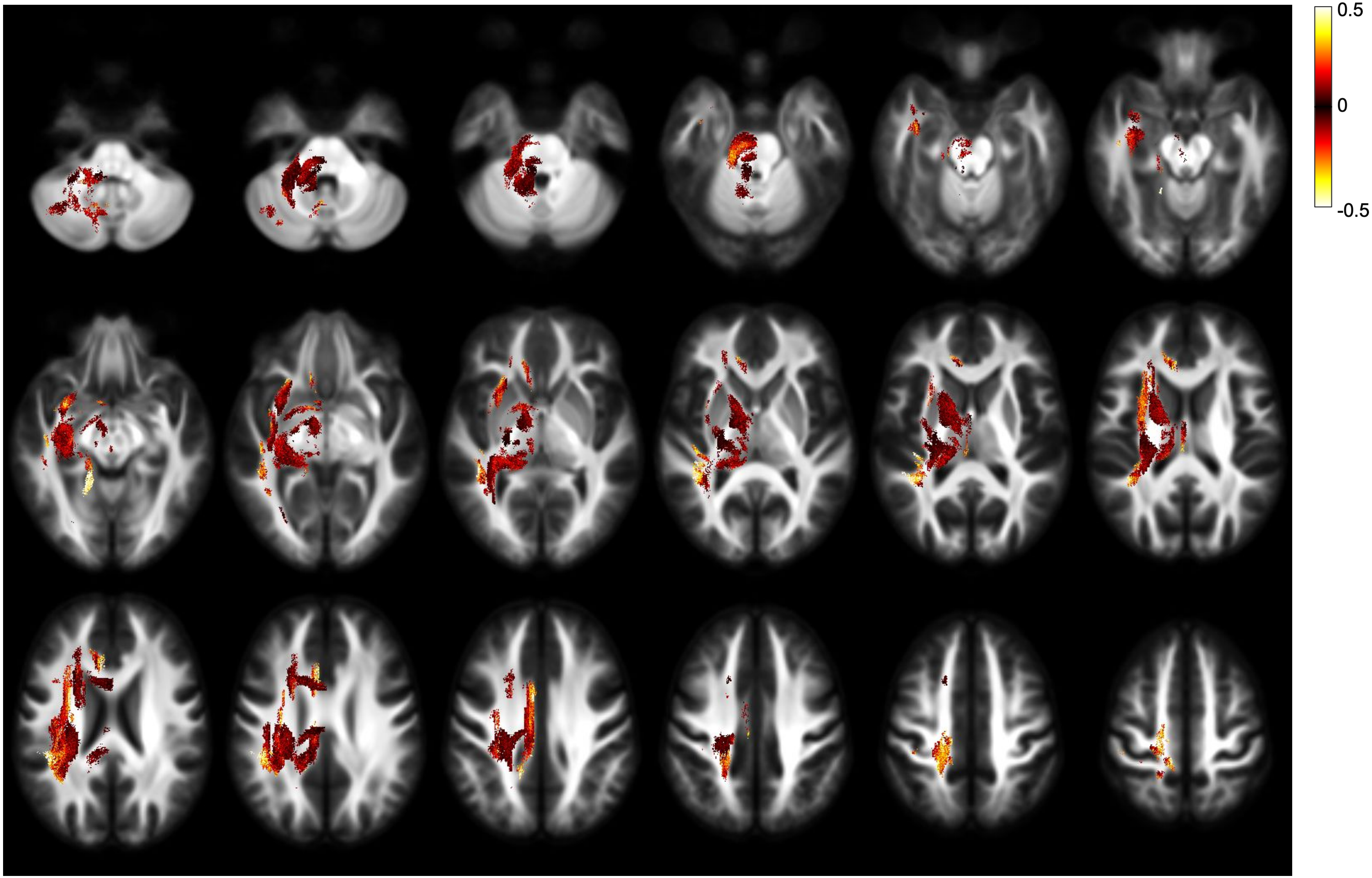


LLD - FDC - effect size - both directions

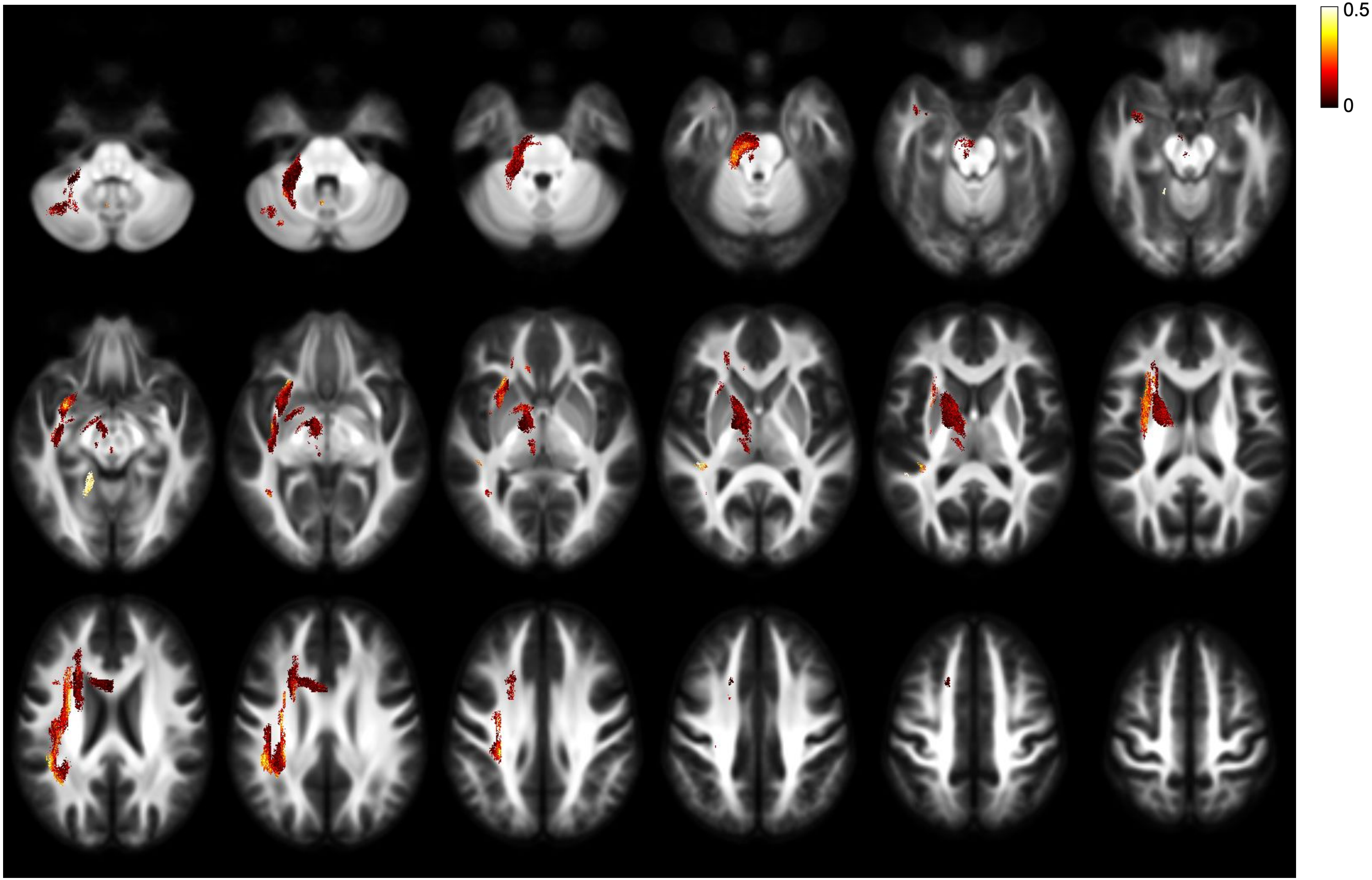


LLD - FDC - effect size - right lateralisation

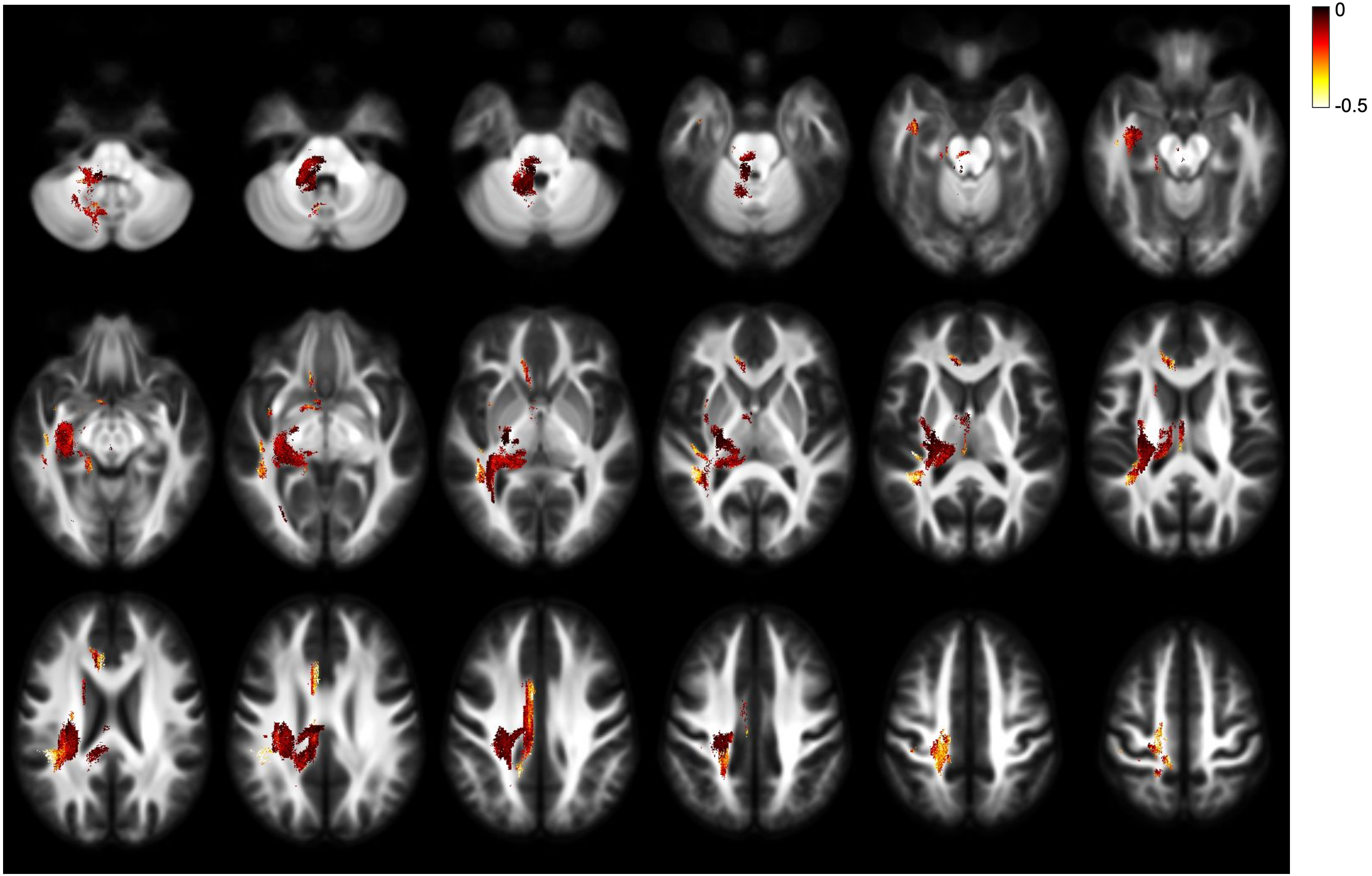


LLD - FDC - effect size - left lateralisation

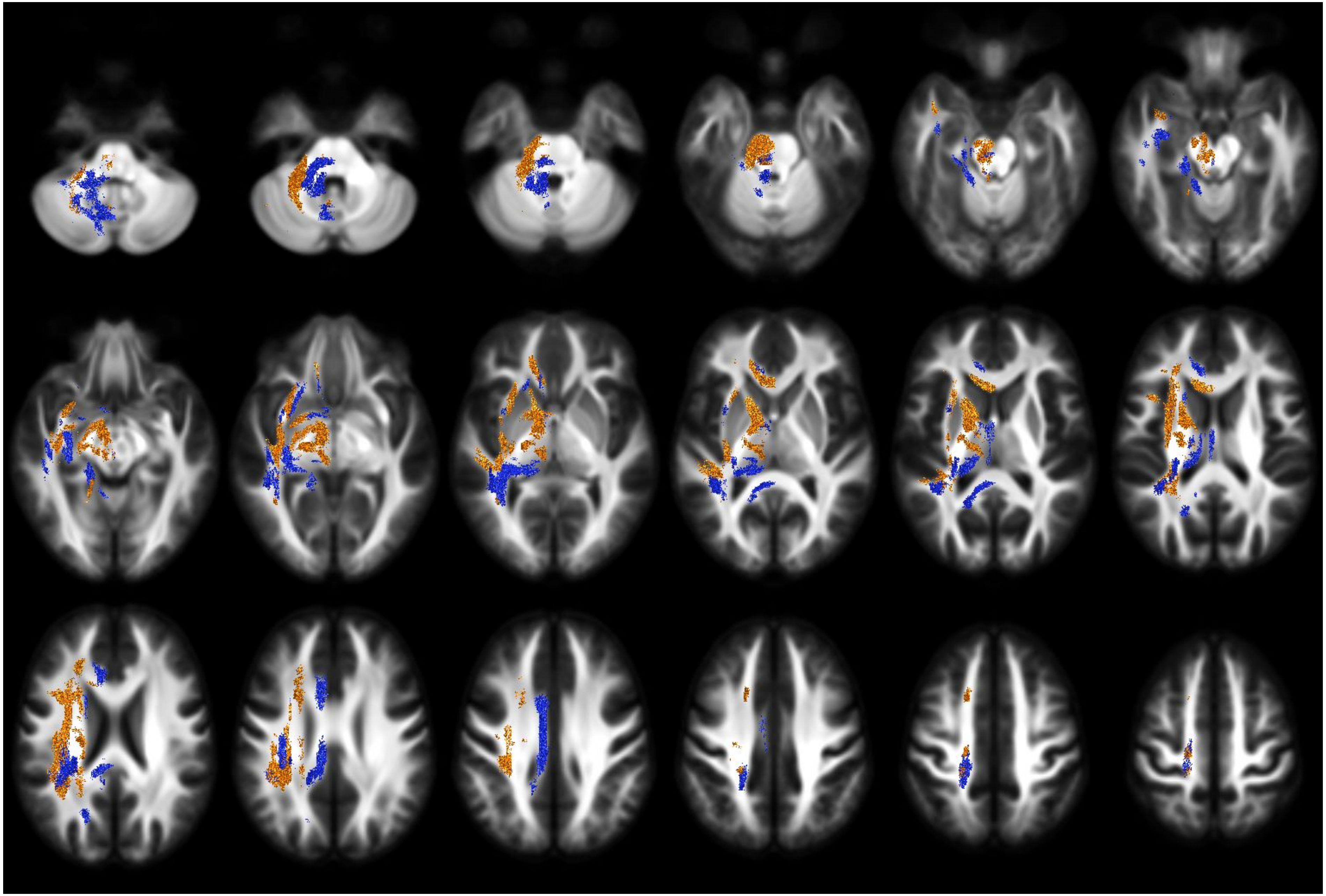

LLD - FD - direction of lateralisation

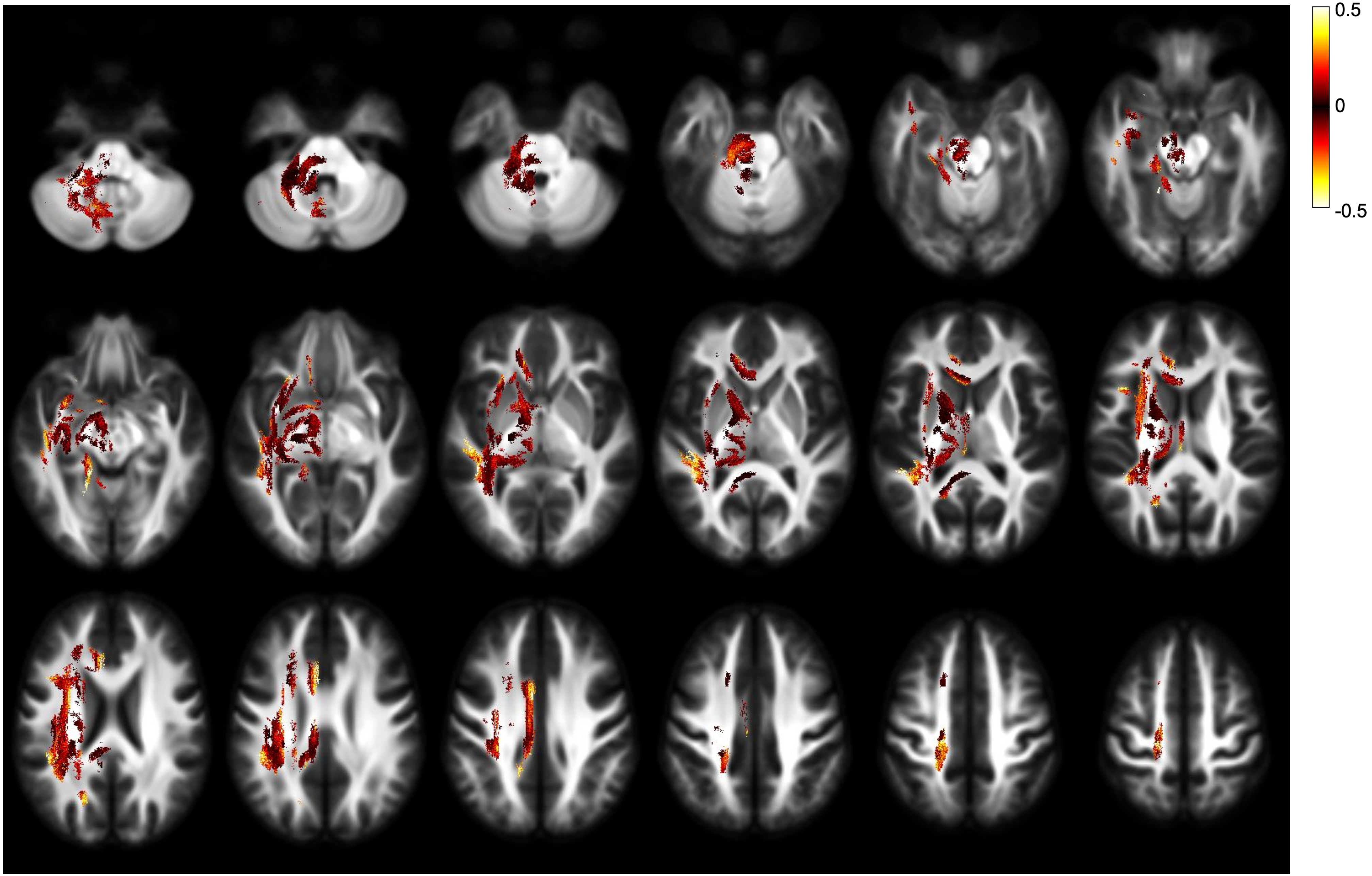


LLD - FD - effect size - both directions

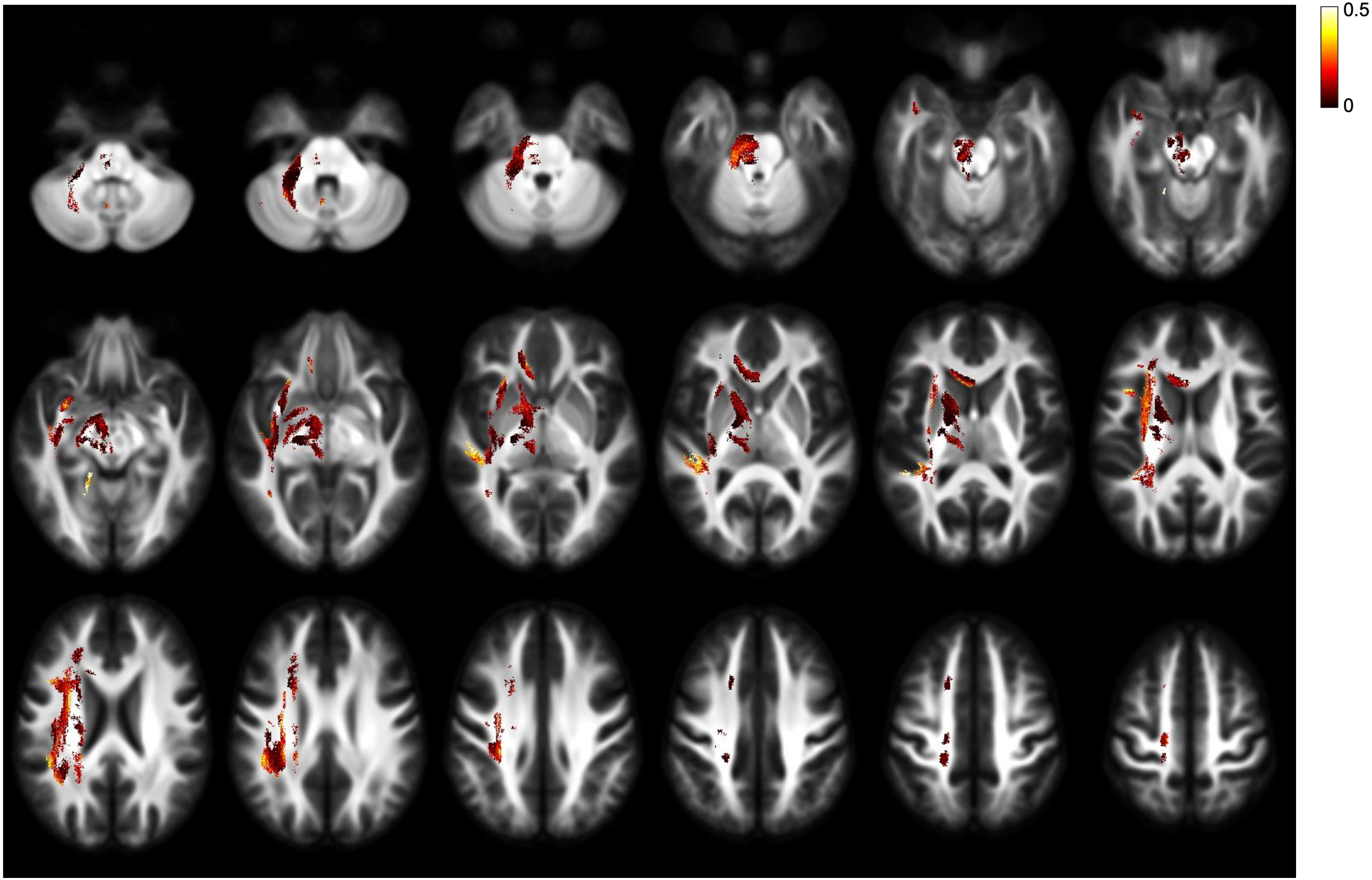


LLD - FD - effect size - right lateralisation

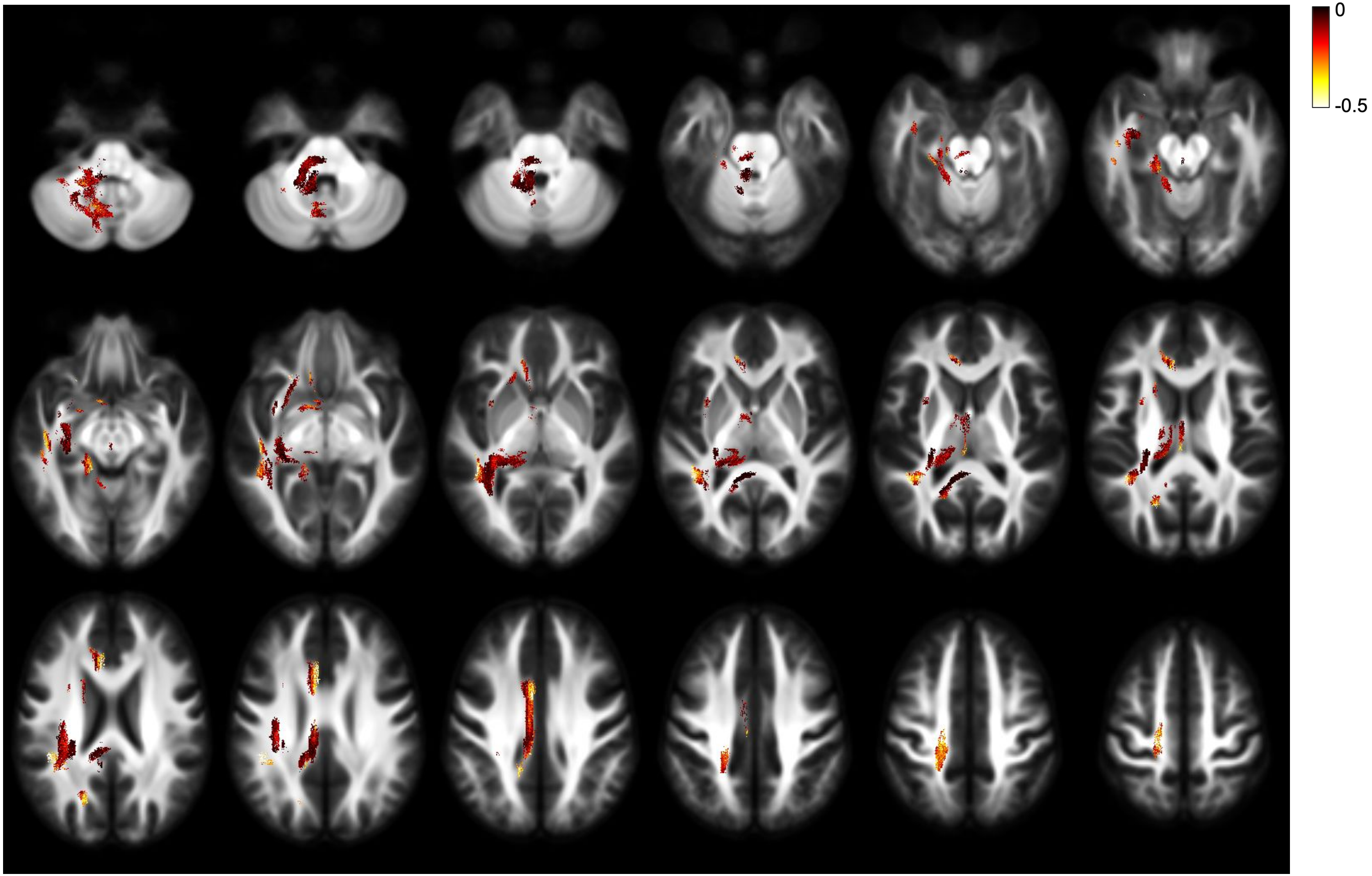


LLD - FD - effect size - left lateralisation

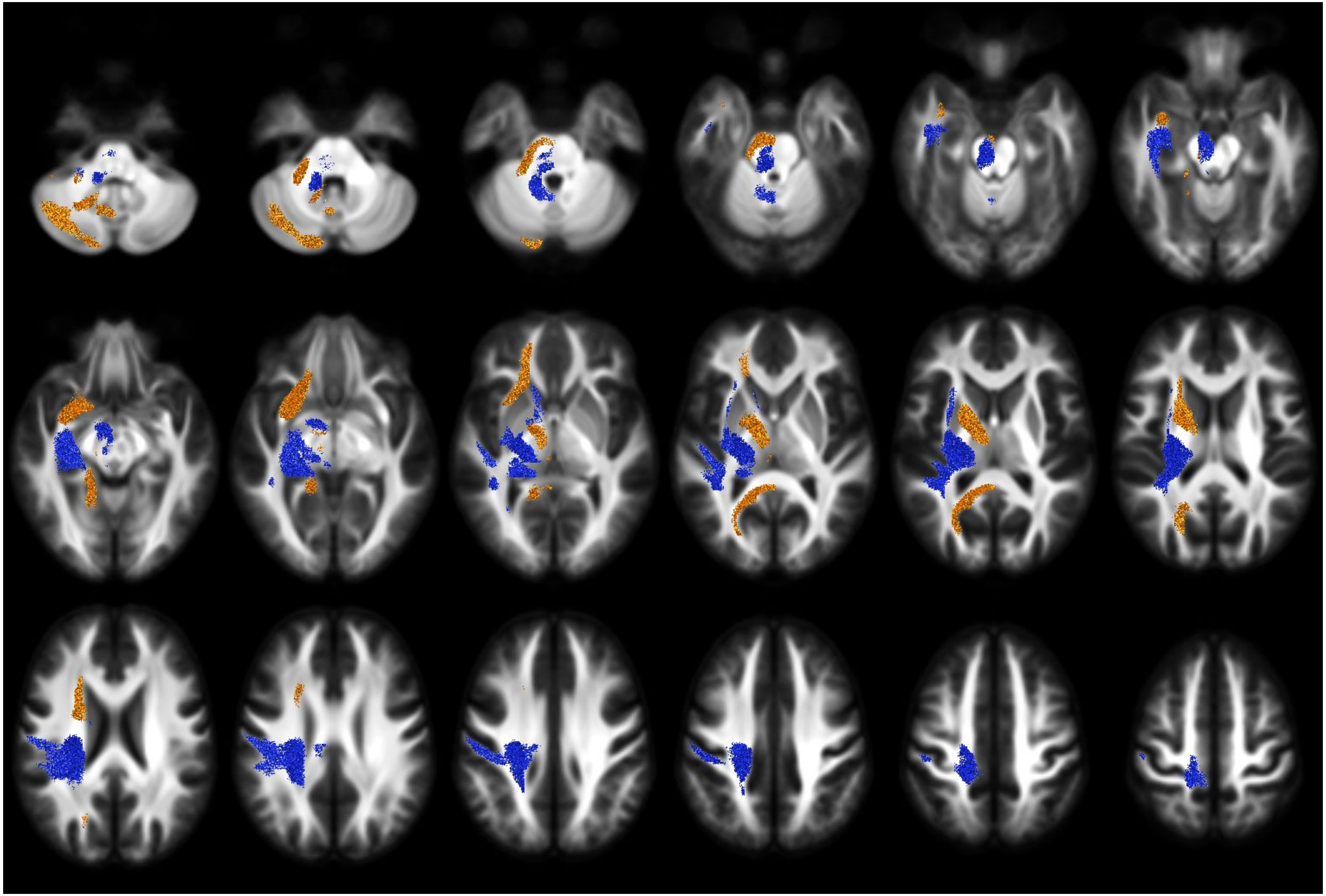

LLD - FC - direction of lateralisation

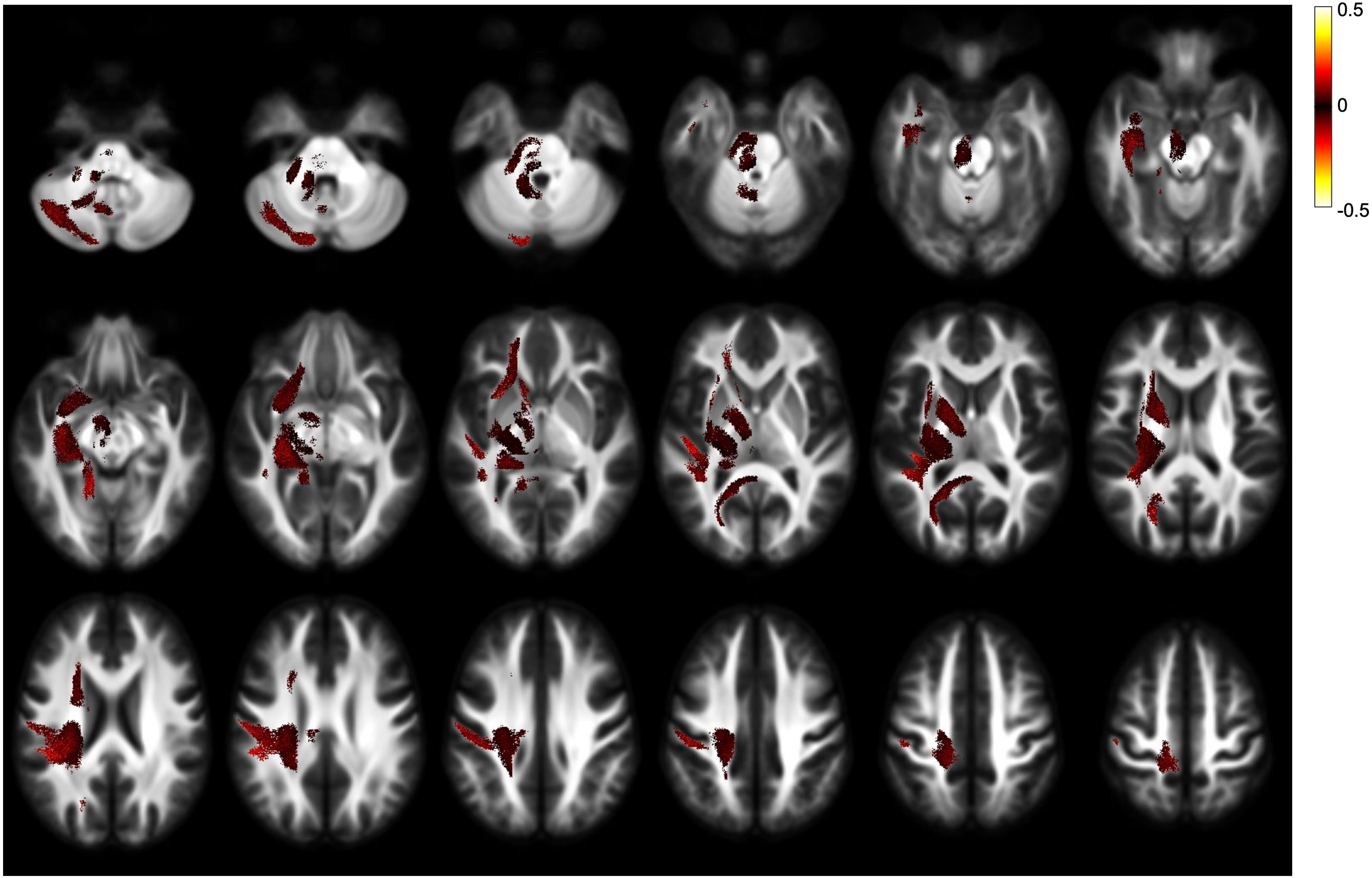


LLD - FC - effect size - both directions

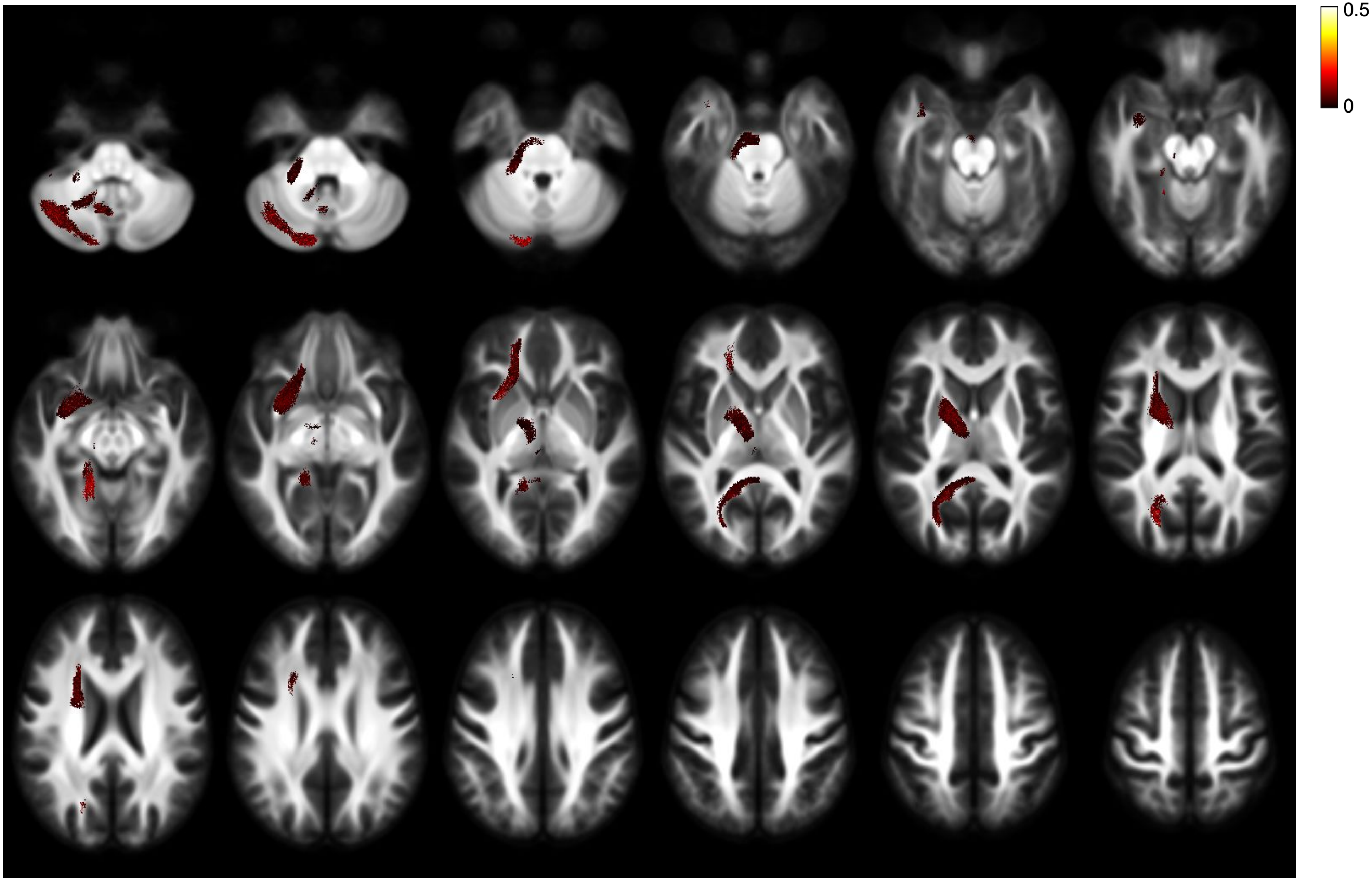


LLD - FC - effect size - right lateralisation

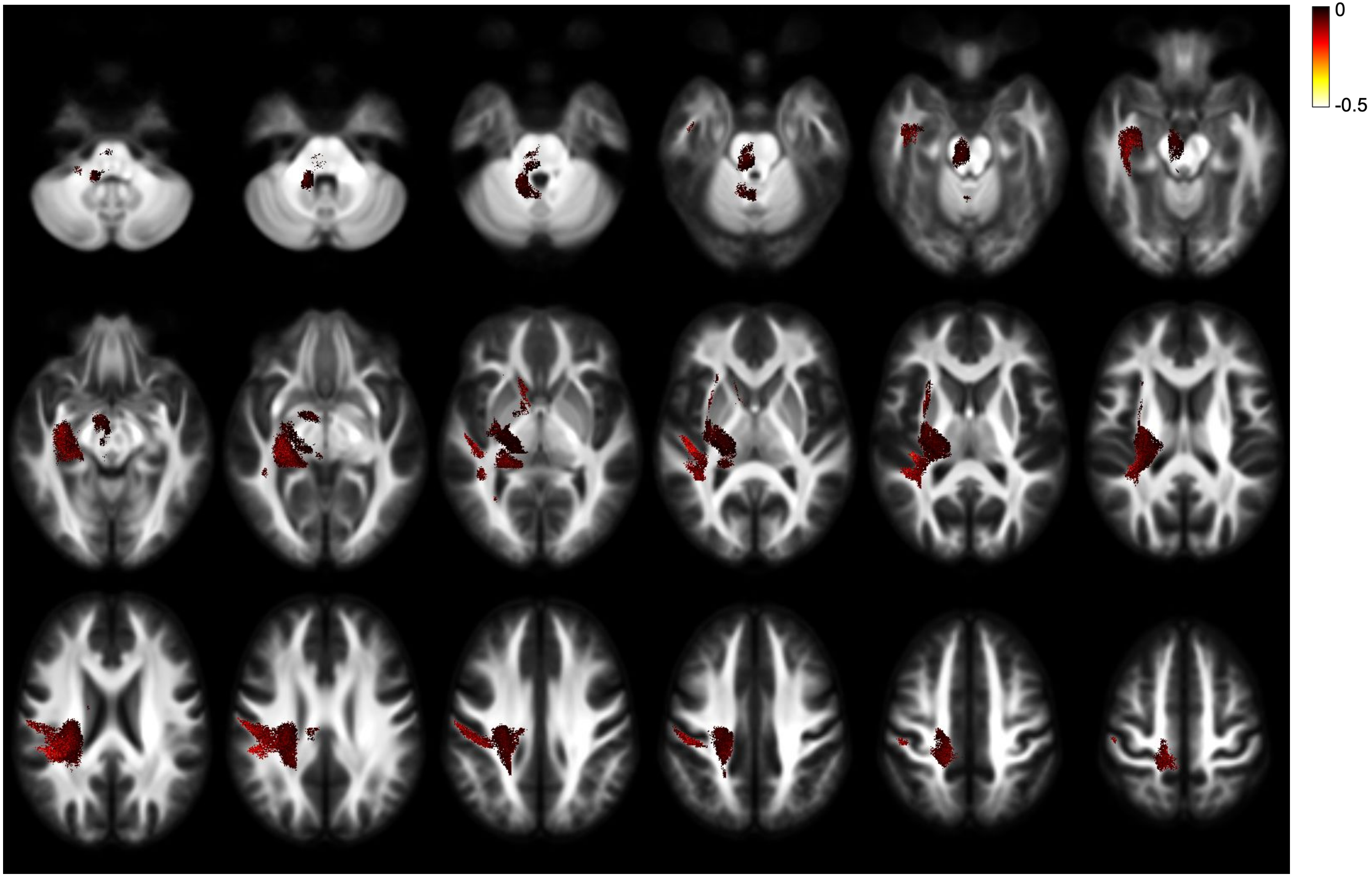


LLD - FC - effect size - left lateralisation

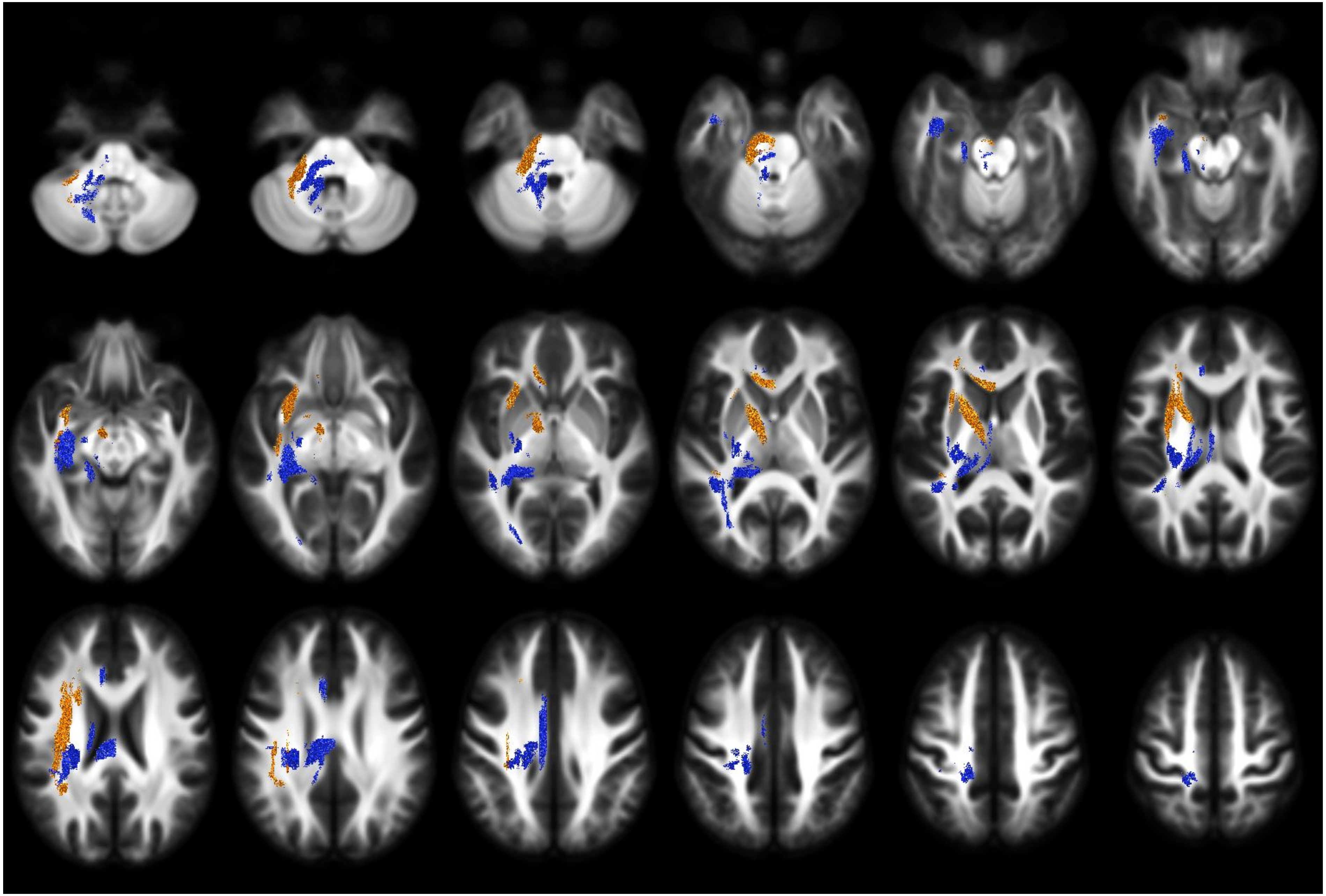

RLD - FDC - direction of lateralisation

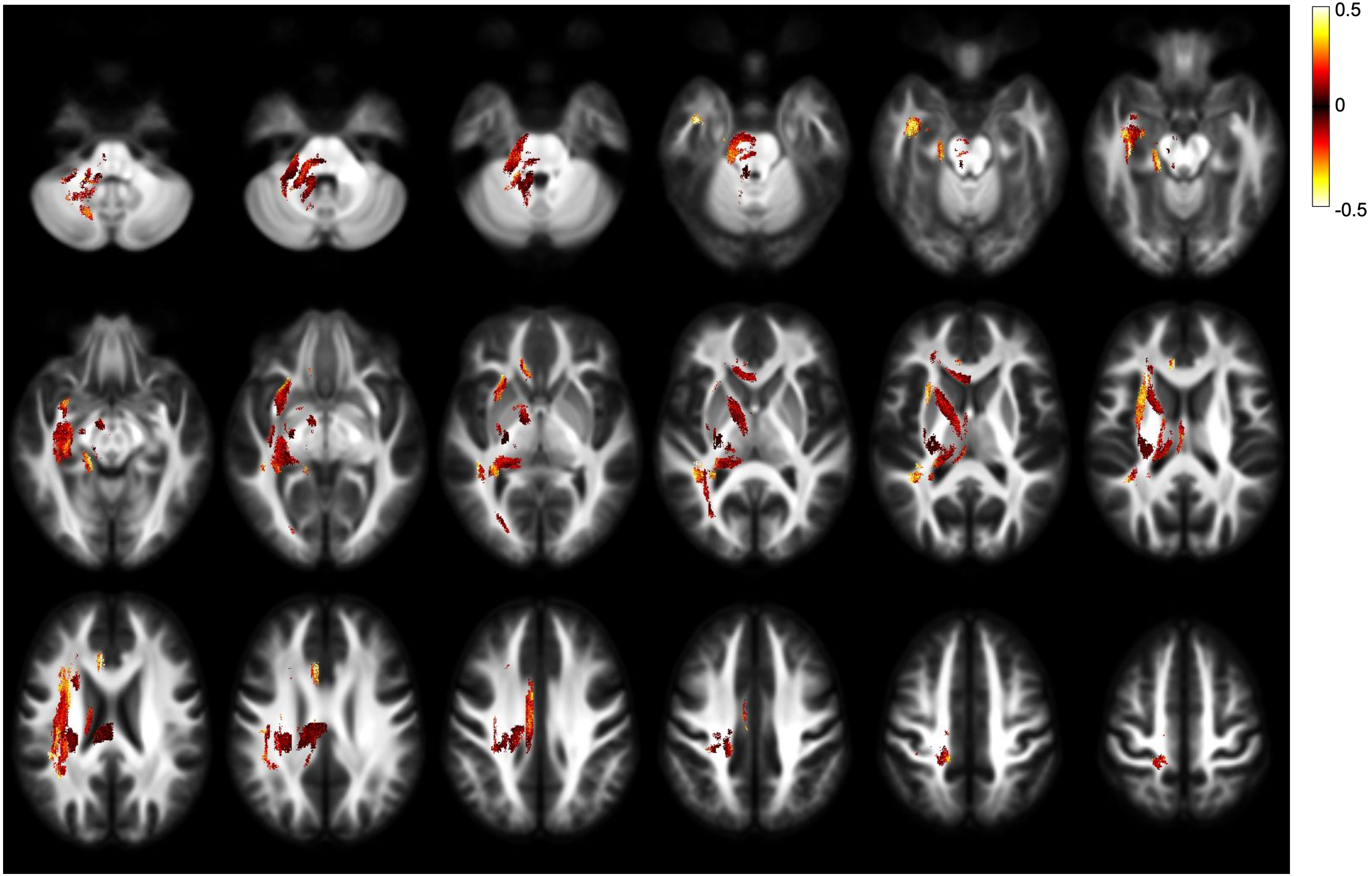


RLD - FDC - effect size - both directions

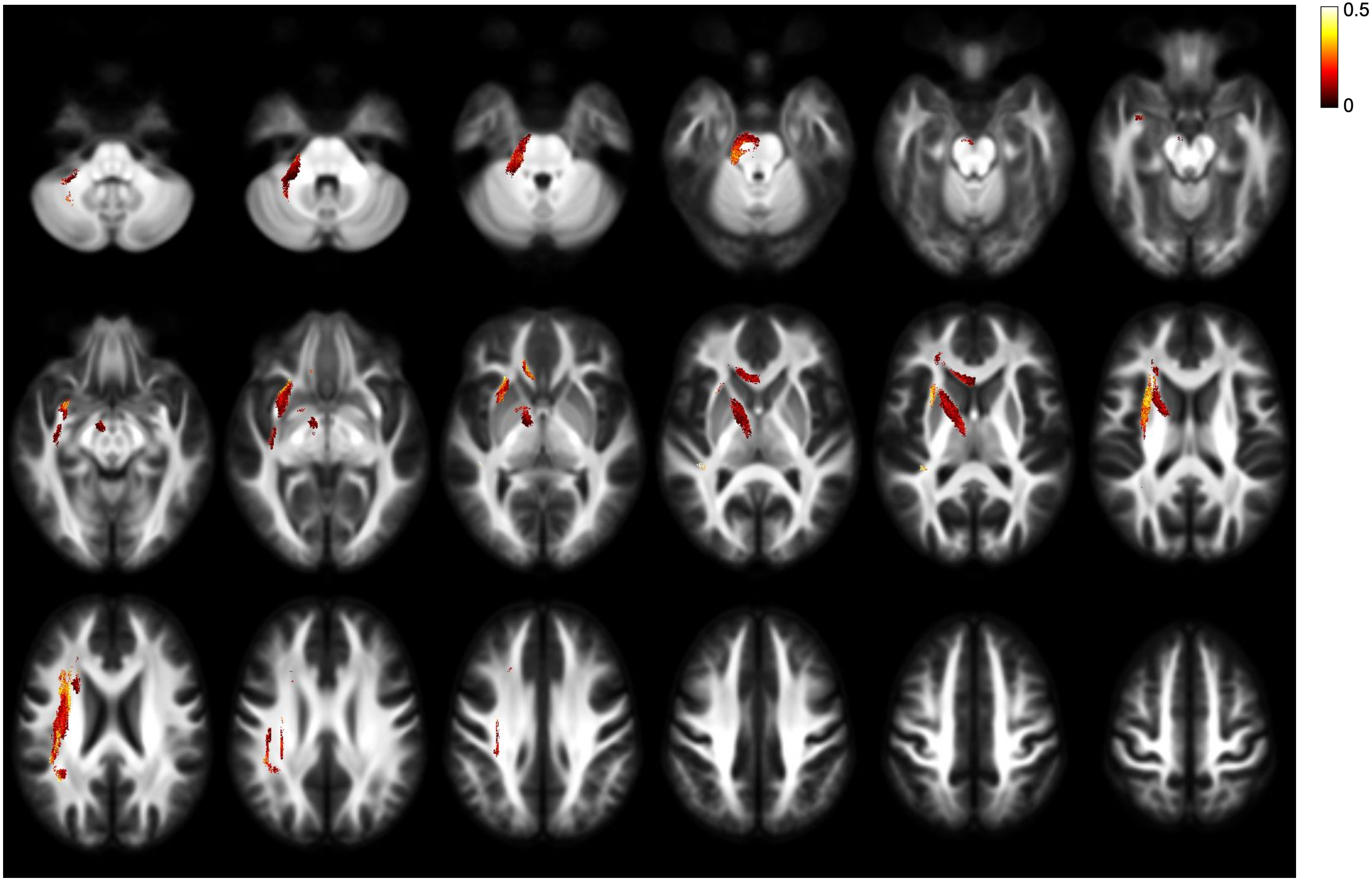


RLD - FDC - effect size - right lateralisation

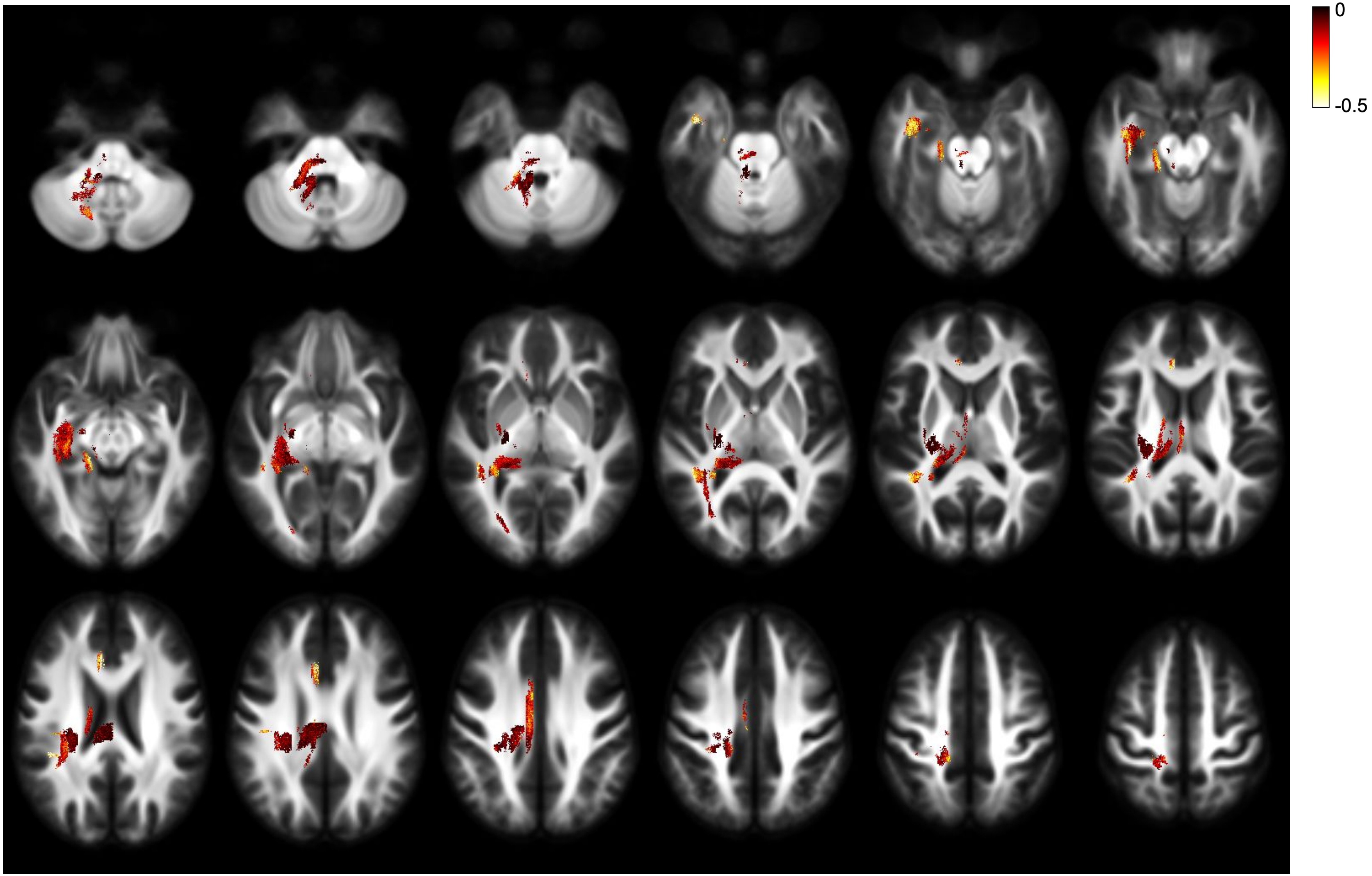


RLD - FDC - effect size - left lateralisation

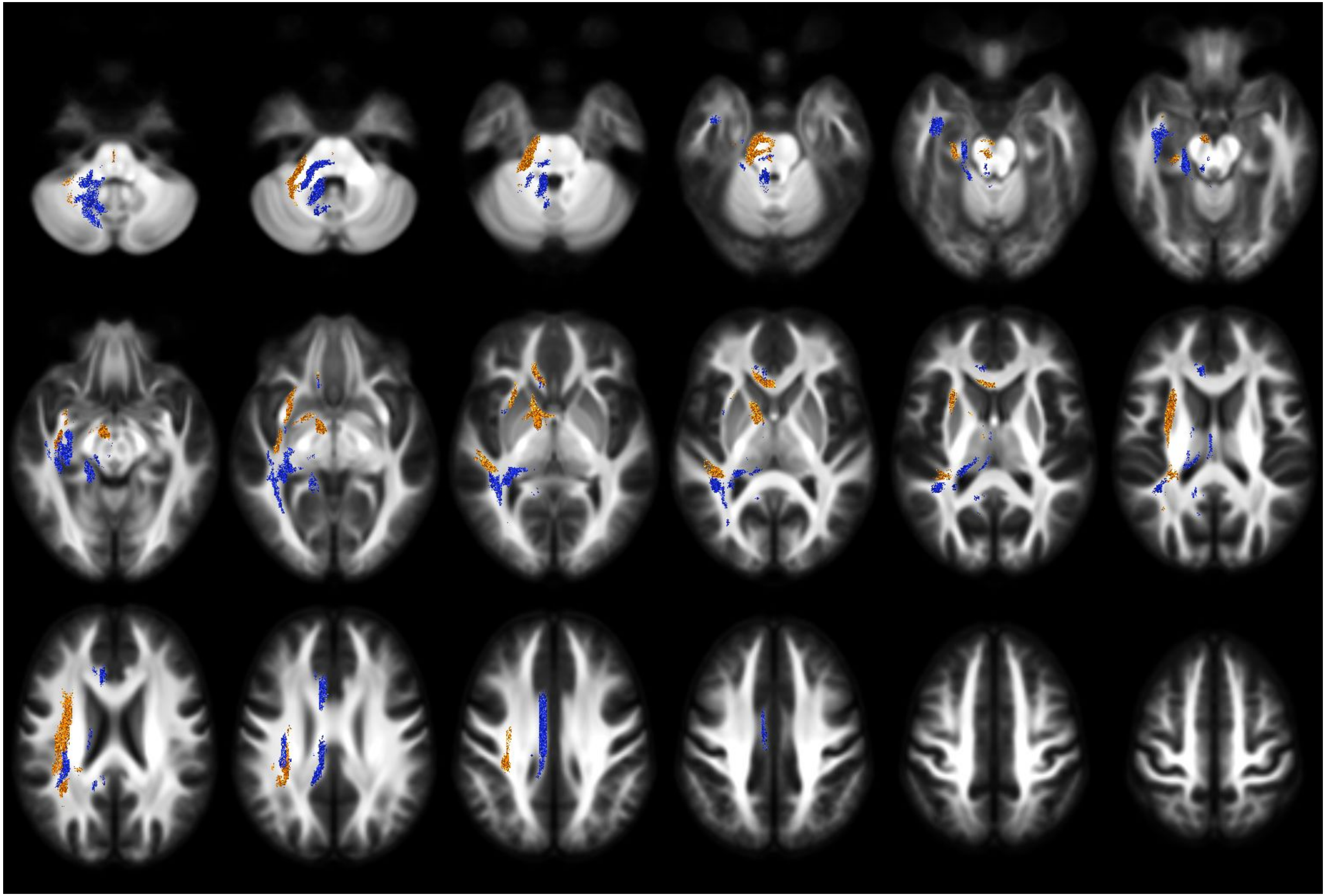

RLD - FD - direction of lateralisation

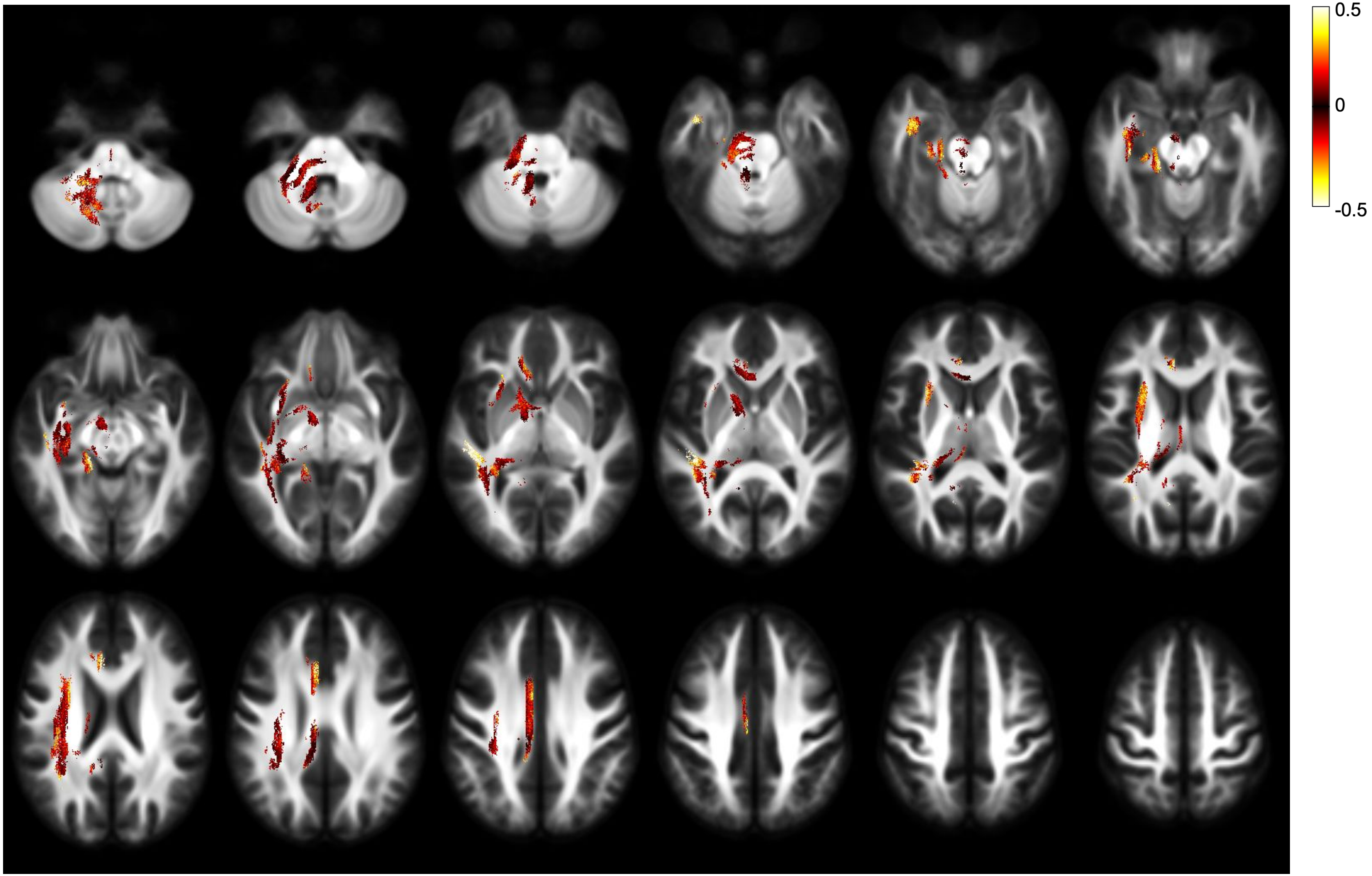


RLD - FD - effect size - both directions

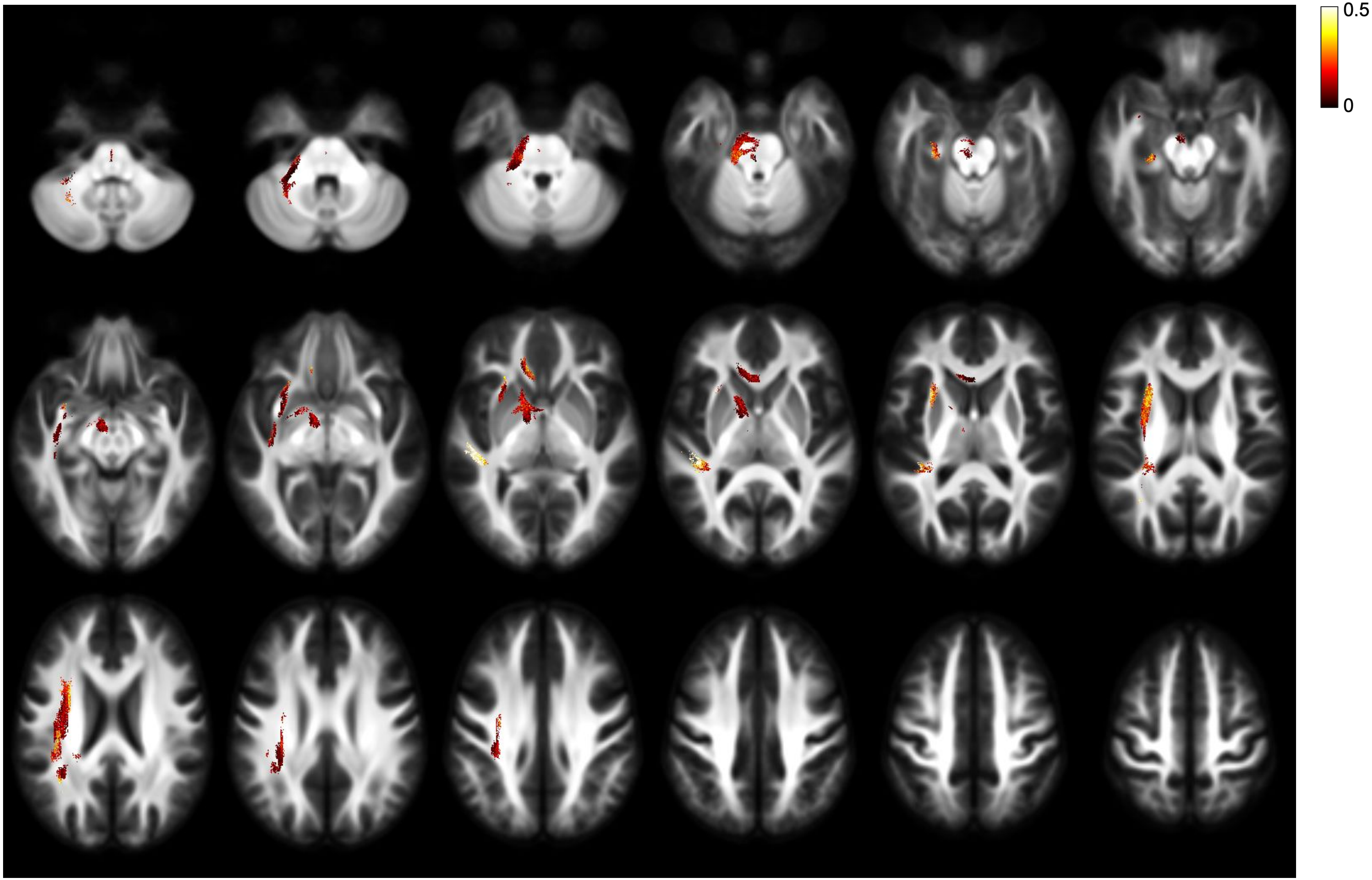


RLD - FD - effect size - right lateralisation

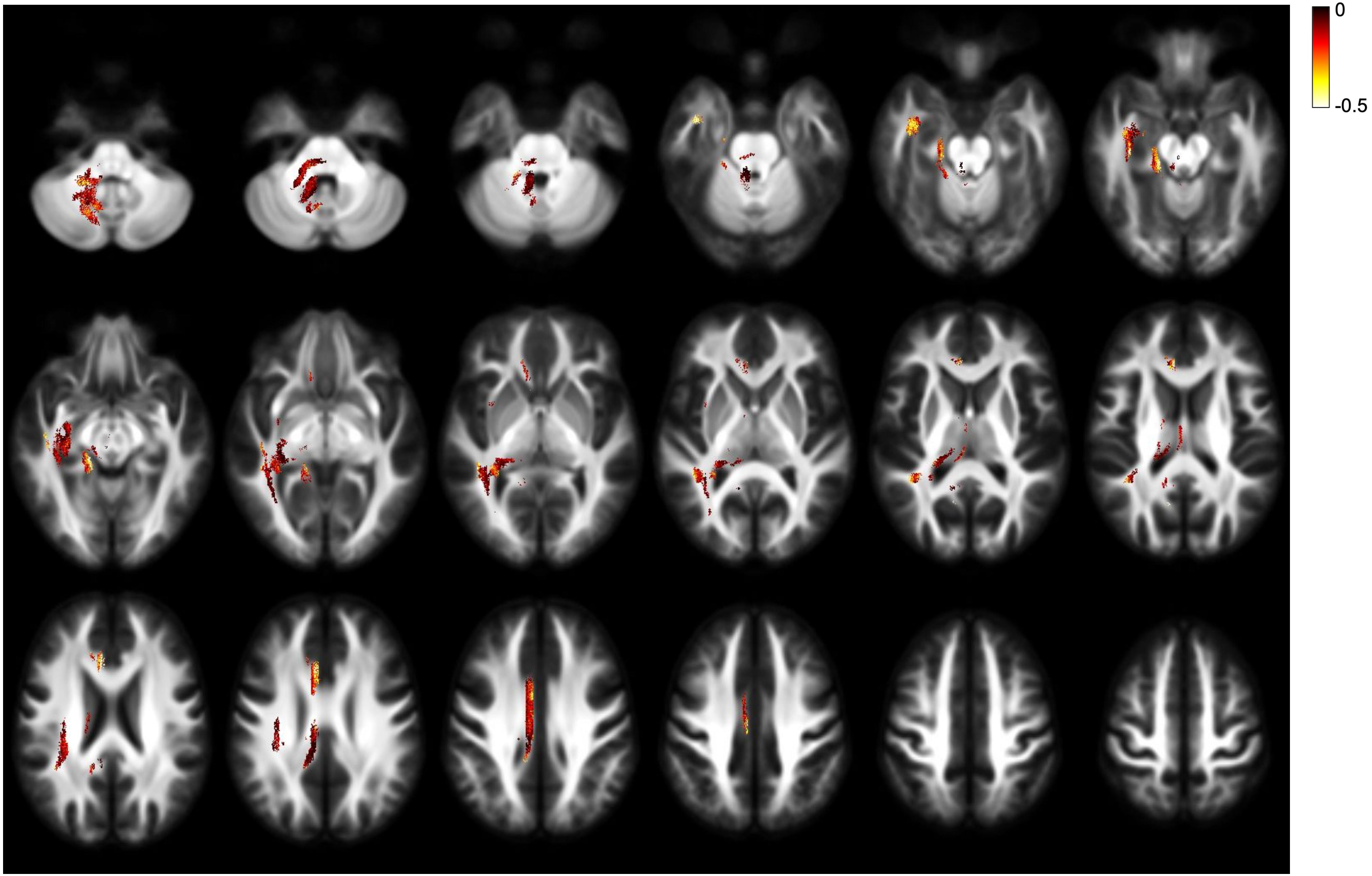


RLD - FD - effect size - left lateralisation

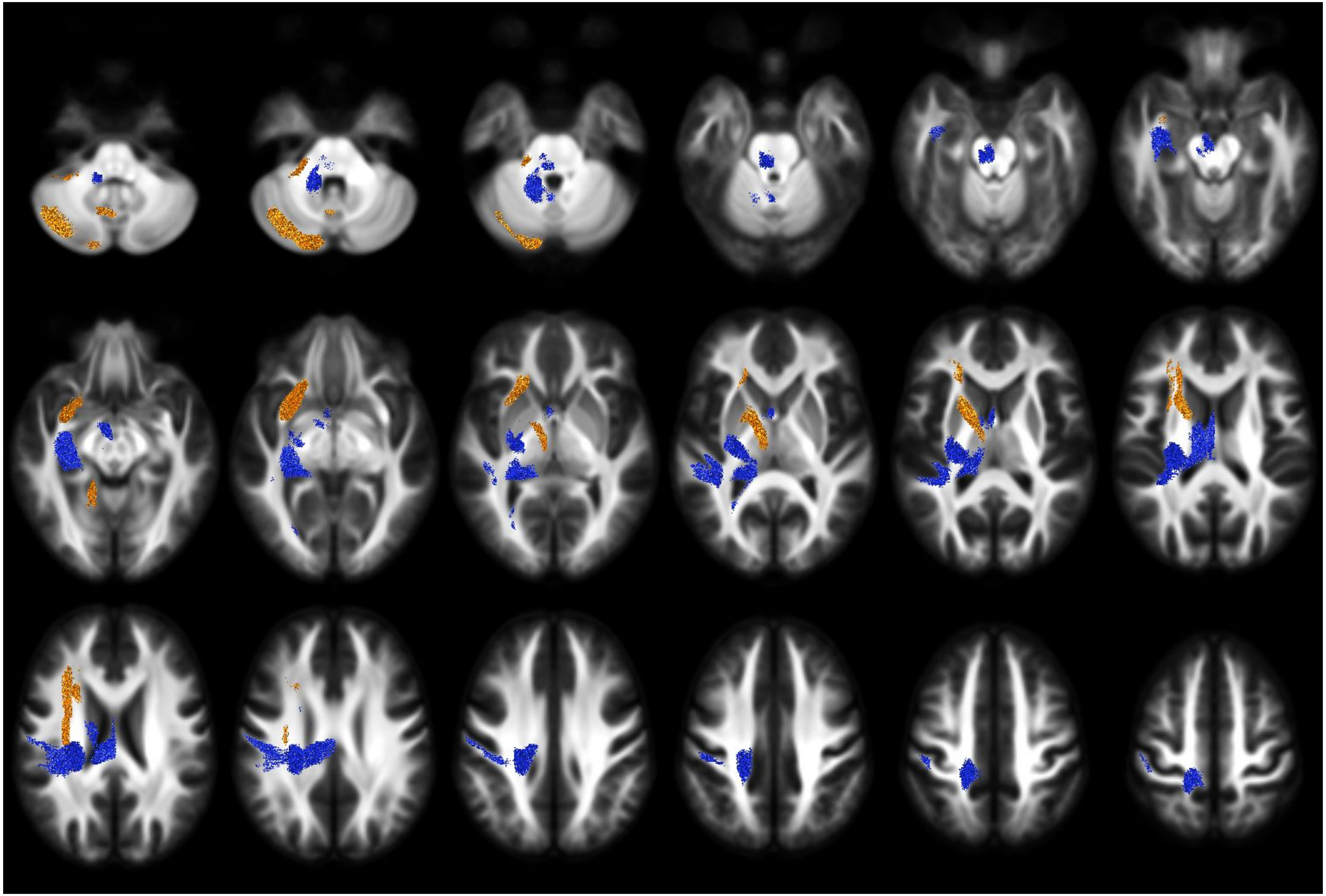

RLD - FC - direction of lateralisation

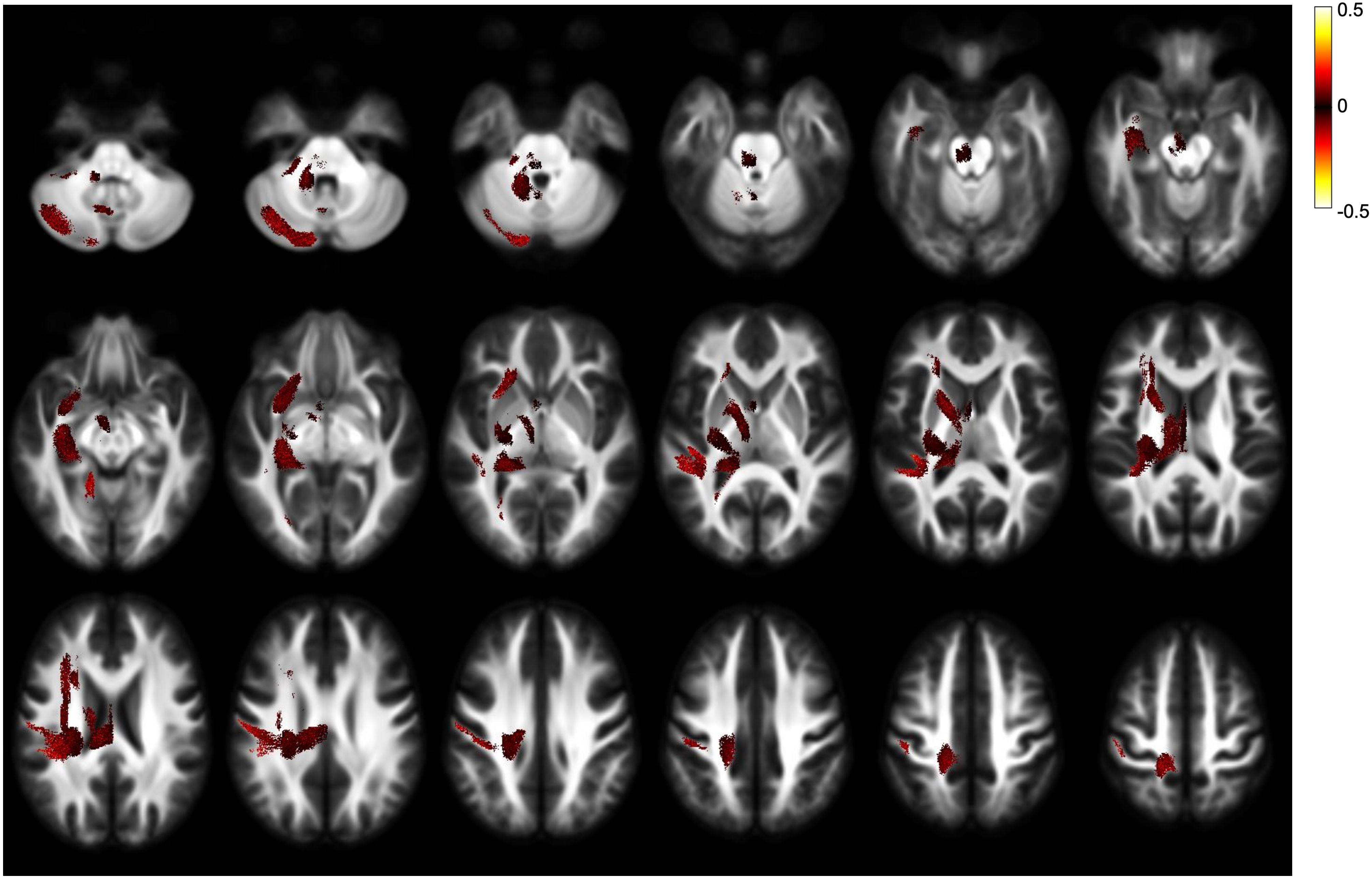


RLD - FC - effect size - both directions

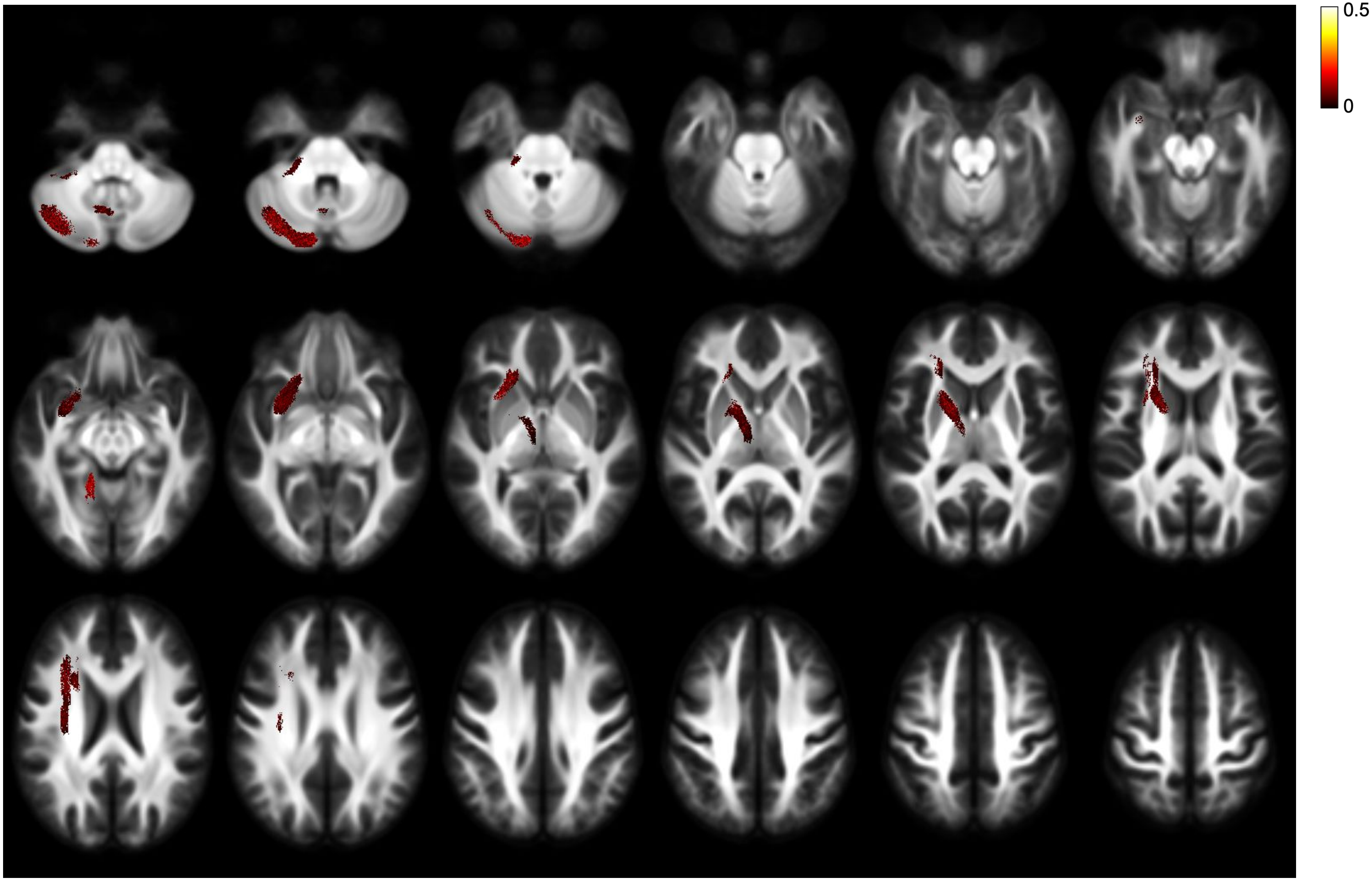


RLD - FC - effect size - right lateralisation

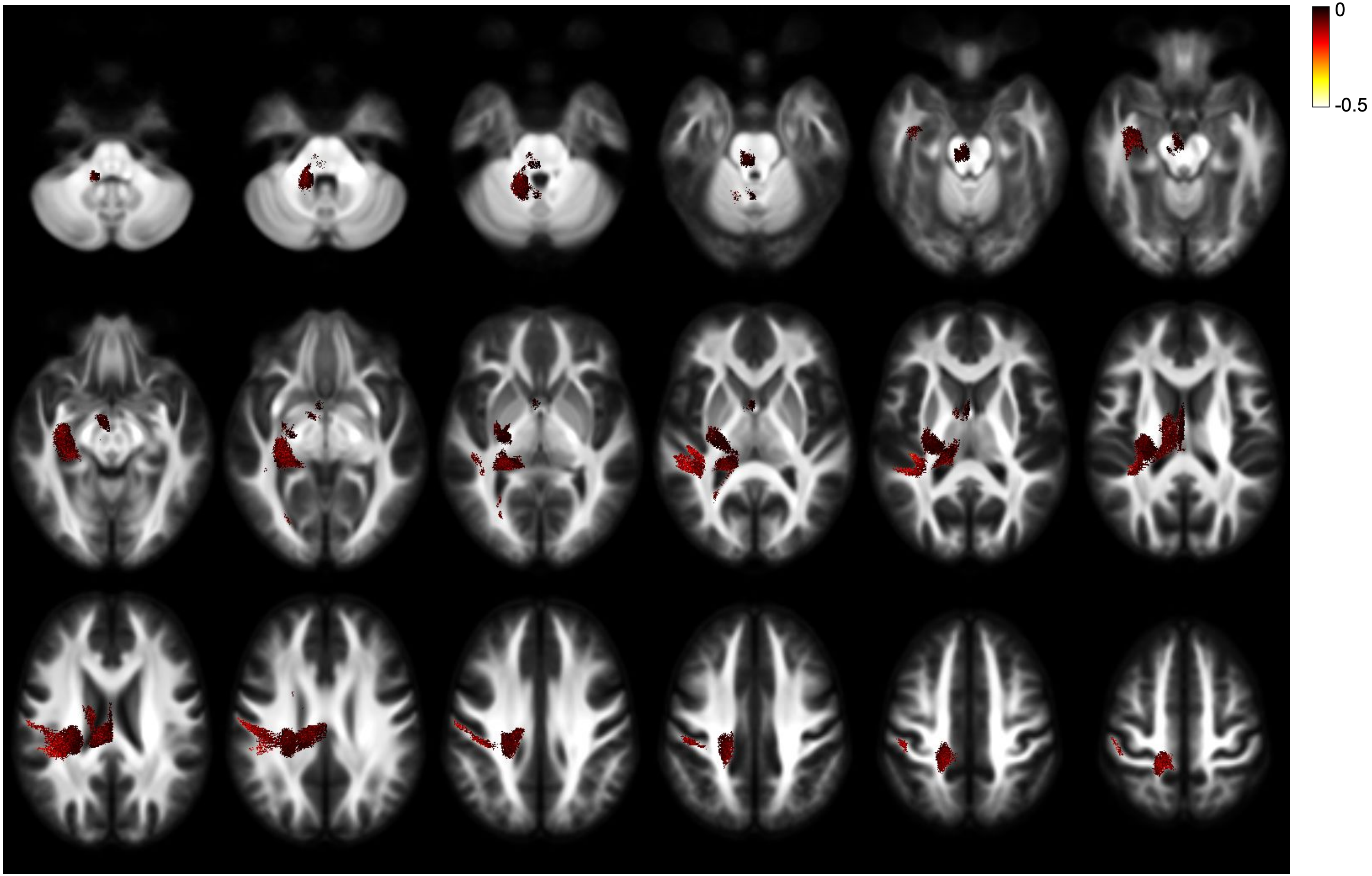


RLD - FC - effect size - left lateralisation
